# Mechanisms controlling the plasma membrane targeting and the nanodomain organization of the plant SPFH protein HIR2

**DOI:** 10.1101/2025.04.10.648198

**Authors:** Michal Daněk, Omar Hdedeh, Jessica Boutet, Héla Safi, Anas Abuzeineh, Amanda Martín-Barranco, Jean-Bernard Fiche, Caroline Mercier, Marcelo Nollmann, Yohann Boutté, Véronique Santoni, Sébastien Mongrand, Alexandre Martinière, Enric Zelazny

## Abstract

HIR2 is a plant-specific protein belonging to the superfamily of SPFH domain-containing proteins that were proposed to play scaffolding functions in membranes. HIR2 organizes in plasma membrane (PM) nanodomains that correspond to nanometric scale structures enriched in specific lipids and proteins acting as signaling/regulation hubs. So far, how PM nanodomains are formed and maintained in plant cells remains largely unknown. Combining state of the art microscopy techniques, we investigated the mechanisms governing the trafficking and the organization into nanodomains of Arabidopsis HIR2 protein. We revealed that the mono S-acylation of HIR2 either on C6 or on C7 was required for HIR2 targeting to the PM of Arabidopsis cells, independently of the conventional secretory pathway. Investigating the mechanisms implicated in the arrangement of HIR2 into nanodomains, we provided evidences that the lipid composition in sterols and very long chain fatty acids of the PM influenced HIR2 organization. HIR2 forms oligomers and we demonstrated here that the C-terminal part of HIR2 is required for self-assembly, similarly to animal SPFH proteins. Interestingly, we highlighted that the oligomerization of HIR2 is essential for its organization in nanodomains and to ensure HIR2 lateral stability in the PM. Overall, we propose that HIR2 nanodomain organization is a complex mechanism relying on different parameters including PM lipid composition and oligomerization. HIR proteins are involved in plant immunity. Here, we revealed that HIR2 nanodomain organization is required to boost the apoplastic ROS burst induced by the bacterial peptide flg22.

**One sentence summary:** S-acylation and oligomerization control the plasma membrane targeting and the organization into nanodomains of the Arabidopsis SPFH-domain containing protein HIR2, respectively.

## INTRODUCTION

Plant plasma membranes (PM) contain myriads of diverse domains at nanometer scale size (20 nm to 1 µm) termed nanodomains that correspond to the functional assembly of specific proteins and lipids (Jarsch et al., 2014; Jaillais and Ott, 2020; Martiniere and Zelazny, 2021; Jaillais et al., 2024). Although the molecular function of nanodomains remains largely unknown, they emerge as signaling/regulatory platforms gathering proteins involved in various cellular processes (Demir et al., 2013; Bucherl et al., 2017; Gronnier et al., 2017; Platre et al., 2019; Smokvarska et al., 2020). The Stomatin/Prohibitin/Flotillin/HflK/C (SPFH) superfamily of proteins is a very diverse family of prokaryotic and eukaryotic membrane proteins that carry a conserved N-terminal domain called the SPFH domain (Rivera-Milla et al., 2006; Hinderhofer et al., 2009). The amino acid sequences of SPFH domain-containing proteins display relatively low similarities among subgroups. Although it is still a matter of debate, SPFH proteins were proposed to have emerged independently in different kingdoms, suggesting their important cellular function (Rivera-Milla et al., 2006). SPFH proteins are arranged in nanodomains in the membranes of diverse subcellular compartments including PM, endoplasmic reticulum (ER), Golgi apparatus and mitochondrial membranes (Browman et al., 2007). So far, SPFH proteins were showed to be involved in various mechanisms related to membranes. Among others, Stomatin, Mechanosensory protein-2 and Podocin play a role in the regulation of ion channels in animals (Price et al., 2004; Huber et al., 2006) and Flotillins (Flots) are implicated in specific endocytic mechanisms in both animal and plant cells (Glebov et al., 2006; Li et al., 2012). SPFH proteins have been proposed for a long time to act as scaffolding proteins in membrane nanodomains (Langhorst et al., 2005), although experimental evidences for such a role remain limited. Nevertheless, an interesting example is provided in plants by *Medicago truncatula* FLOT4 protein that acts as a central hub during primary nanodomain assembly by recruiting the remorin SYMREM1 that in turn interacts with and stabilizes the activated LYSINE MOTIF KINASE 3 (LYK3) receptor into nanodomains, insuring symbiotic root infection (Liang et al., 2018). How SPFH proteins assemble in nanodomains is not known but this process may rely on their propensity to form large homo- and hetero-oligomers (Browman et al., 2007). Early observations by electron microscopy highlighted that mitochondrial prohibitins from yeast form ring-shaped structures (Tatsuta et al., 2005). Recently, the structure of SPFH proteins from *Escherichia coli* named HflC and HflK revealed that these proteins assemble in a large circular complex composed of 12 copies of HflK–HflC dimers that delimits a laterally segregated membrane domain of 20 nm in diameter (Ma et al., 2022). A similar super-complex attached to the membrane surface of cell-derived vesicles was observed for human Flot1/Flot2 proteins (Fu and MacKinnon, 2024). These very interesting works strengthen the role of SPFH proteins in nanodomain structuration through a possible compartmentation of lipids and proteins. So far, the domains involved in the oligomerization of SPFH proteins were mostly identified in their C-terminal parts such as for animal and plant Flot, in the so-called Flot domain, and for human Stomatin (Umlauf et al., 2006; Solis et al., 2007; Yu et al., 2017; Singh et al., 2022). However, the structure of the HflK– HflC multimer from *E. coli* revealed complex and multiple interactions between monomers also through their SPFH domains (Ma et al., 2022).

Plants possess a specific group of SPFH proteins named Hypersensitive Induced Reaction (HIR) proteins composed of four members in *Arabidopsis thaliana* (HIR1 to HIR4). HIR contribute to immunity against various pathogens such as bacteria, viruses and fungi in several plant species (Mei et al., 2020; Liu et al., 2024). The mutation of *HIR2* and *HIR3* genes in *Arabidopsis thaliana* and the overexpression of HIR3 protein in rice induce a sensitivity to *Pseudomonas syringae* and a resistance to *Xanthomonas oryzae*, respectively (Qi et al., 2011; Li et al., 2019). HIR proteins from different plant species were found to interact with proteins implicated in the response to pathogens, such as the receptor Flagellin-Sensing 2 (FLS2), reinforcing the link between HIR and plant immunity (Jung and Hwang, 2007; Zhou et al., 2009; Qi et al., 2011; Qi and Katagiri, 2012; Hdedeh et al., 2025). Although the underlying molecular mechanisms involving HIR proteins remain enigmatic, HIR are implicated in the reactive oxygen species (ROS) burst occurring in response to the bacterial peptide flg22 (Liu et al., 2024). HIR proteins do not contain transmembrane domains, but are associated with membranes (Danek et al., 2016). Except for HIR3, all the Arabidopsis HIR proteins were shown to arrange in nanodomains in the PM and display low lateral mobility dictated in part by the cell wall (Danek et al., 2020). On the other hand, biotic and abiotic stresses were shown to affect Arabidopsis HIR1 nanodomain dynamics (Xu et al., 2022). In animal cells, Flot2 association with membranes is ensured by myristoylation and S-acylation (Neumann-Giesen et al., 2004). Myristoylation and S-acylation are post-translational lipidation modifications of a target protein that consist in the addition of a myristate to a N-terminal glycine through an amide bond and the addition of a fatty acyl group to cysteine residues via a thioester bond, respectively (Hemsley, 2015). When a palmitate is linked to a protein, S-acylation can be referred to as palmitoylation. Contrary to myristoylation, S-acylation is reversible which confers to this lipid modification the potential to regulate protein function in response to stimuli (Hurst and Hemsley, 2015). Myristoylation and S-acylation are well known to provide hydrophobic anchors allowing the association of proteins to membranes in plant cells (Hemsley, 2015). The S-acylation of some *Arabidopsis thaliana* and *Medicago truncatula* remorins contributes to their PM association but is not responsible for their arrangement in nanodomains (Konrad et al., 2014). S-acylation function is not restricted to membrane association since S-acylation can be also implicated in the formation and stabilization of protein complexes (Lakkaraju et al., 2012; Hurst et al., 2023). Interestingly, Arabidopsis HIR proteins were shown to be myristoylated and S-acylated (Hemsley et al., 2013; Majeran et al., 2018; Kumar et al., 2022). Similarly to other SPFH proteins, HIR are able to homo- and hetero-oligomerize (Qi et al., 2011; Liu et al., 2024). However, it is not known whether the oligomerization plays a role in the recruitment of HIR into membrane nanodomains or in the organization of the domains themselves.

In plants, the lipid composition of the PM influences the dynamics of proteins and their organization into nanodomains (Gronnier et al., 2018; Jaillais and Ott, 2020). Among lipids, the role of sterols was probably the most explored since sterols, together with sphingolipids, were historically described to be important for nanodomain organization by forming liquid ordered domains in the PM (Grosjean et al., 2015; Mamode Cassim et al., 2019). Arabidopsis plants treated with the sterol-depleting agent methyl-β-cyclodextrin (mβCD) or the sterol biosynthesis mutant *cyclopropylsterol isomerase 1 (cpi1-1)* both display altered intracellular dynamics of Flot1 (Li et al., 2012; Cao et al., 2020). Interestingly, both sterols and phosphatidylinositol 4-phosphate are required for the nanodomain organization of *Solanum tuberosum* StREM1;3 in the PM (Gronnier et al., 2017). Phosphatidylserine (PS) is required to stabilize the GTPase Rho-of-Plant 6 (ROP6) in PM nanodomains (Platre et al., 2019). Interestingly, PS and sphingolipids that are both closely linked to the organization of PM nanodomains in plants display a common trait, a very long chain fatty acid (VLCFA) that correspond to fatty acids with more than 18 carbon atoms. These data illustrate that one lipid category can influence the patterning in the PM of functionally unrelated proteins and that multiple lipids species can be required for the proper organization in the PM of a given protein.

Although SPFH proteins were identified more than twenty-five years ago, their function and regulation remain largely unknown in plants. Here, we sought to better understand the mechanisms governing the PM targeting and the organization into nanodomains of the plant-specific HIR proteins, using Arabidopsis HIR2 isoform as a model. We highlighted that the mono S-acylation of HIR2 either on cysteine 6 or on cysteine 7 is mandatory for its PM targeting in Arabidopsis. Combining state of the art microscopy techniques, we showed that: (i) the lipid composition in sterols and VLCFA of the PM influences the organization of HIR2 into nanodomains and HIR2 molecule dynamics in the PM; (ii) HIR2 nano-organization in the PM is driven by HIR2 oligomerization through its C-terminal part. In addition, we provided evidences that the nanodomain-organization of HIR2 is required to stimulate the ROS burst in plant cells in response to flg22.

## RESULTS

### HIR2 and remorins are arranged in distinct nanodomains in the plasma membrane

The PM localization of HIR2 protein in plant cells was previously reported in *Arabidopsis thaliana* and in tobacco leaves by expressing a translational HIR2 fusion with a yellow fluorescent protein (HIR2-YFP) under the control of the strong *35S* constitutive promoter (Qi et al., 2011; Danek et al., 2020). Because overexpression may affect the subcellular localization of a protein and to determine the expression profile of HIR2 protein in *Arabidopsis thaliana*, we generated transgenic lines expressing HIR2 fused to the fluorescent protein mCherry, under the control of *HIR2* promoter (pHIR2::HIR2-mCherry). In the root, *HIR2* promoter drives HIR2-mCherry expression specifically in epidermal cells, in root hairs as well as in the lateral root cap (Supplemental Figure S1). HIR2-mCherry fluorescence was also observed in hypocotyl cells and in cotyledon pavement cells. At the subcellular level, whatever the imaged tissue, HIR2-mCherry protein was only localized in the PM (Supplemental Figure S1), in accordance with previous works (Qi et al., 2011; Danek et al., 2020). We verified the expression of HIR2-mCherry fusion protein in Arabidopsis transgenic lines by performing an anti-mCherry immunoblot. We observed a strong signal at the expected molecular weight of HIR2-mCherry as well as a faint signal of higher molecular weight, both of which being not observed in protein extracts from wild-type (WT) plants, showing the specificity of these signals (Supplemental Figure S2). Similar signals were detected when HIR2-mCherry was transiently expressed under the control of *HIR2* promoter in *Nicotiana benthamiana* leaves, whereas no signal was detected in uninfiltrated leaves (Supplemental Figure S2).

Remorins are abundant PM proteins that are probably the most studied proteins arranging into nanodomains in plants (Raffaele et al., 2009; Jarsch et al., 2014; Gronnier et al., 2017). We investigated the level of co-localization between two remorin isoforms from *Arabidopsis thaliana*, REM1.2 and REM1.3, fused to the YFP (Jarsch et al., 2014), and HIR2-mCherry, the three proteins being expressed under the control of their respective endogenous promoters in Arabidopsis. Spinning-disk microscopy analysis performed on root epidermal cells revealed a very weak co-localization between the two remorins and HIR2 in the PM (Figure 1A, B) demonstrated by low Pearson’s correlation coefficients (PCC) (Figure 1C). These results suggest that HIR2 and remorins are mainly organized in distinct PM nanodomains.

**Figure 1.**
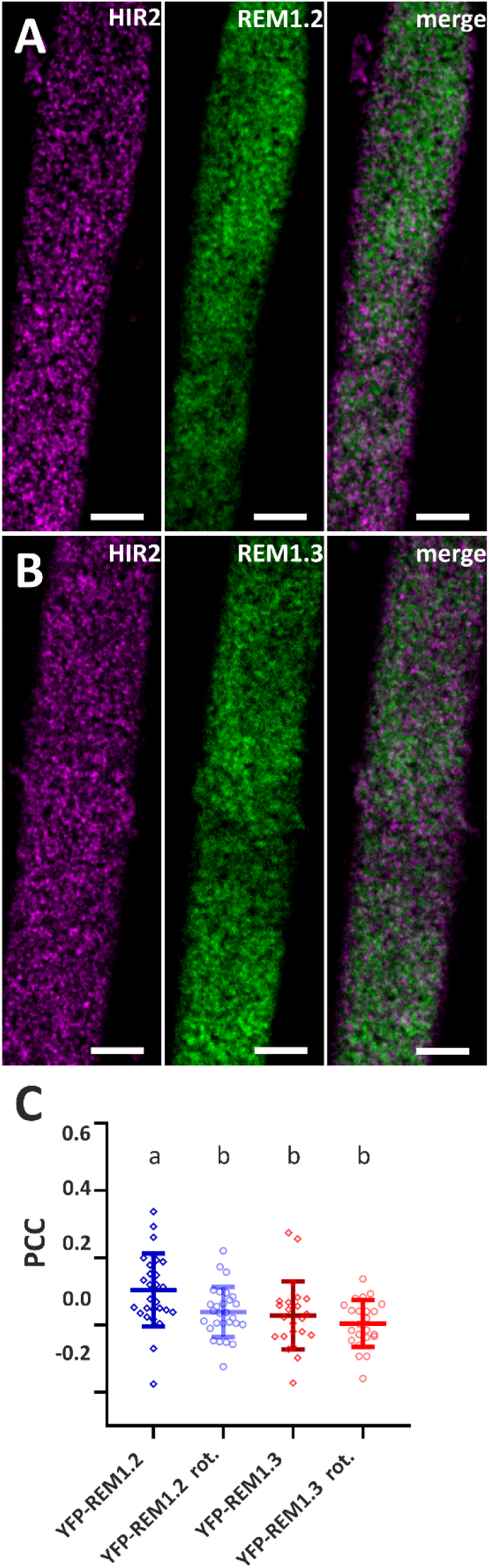
HIR2 and remorins arrange in distinct plasma membrane nanodomains. Spinning disk microscopy analyses of root epidermal cells from pHIR2::HIR2-mCherry Arabidopsis transgenic lines co-expressing pREM1.2::YFP-REM1.2 **(A)** or pREM1.3::YFP-REM1.3 **(B)**. In majority, in (A) and (B), HIR2-mCherry PM nanodomains (left panels) and YFP-REM PM nanodomains (middle panel) do not co-localize (merge, right panel). Arabidopsis seedlings were grown for 11 days on MS/2 medium. Scale bars: 5 µm. **(C)** Quantification of the co-localization in PM nanodomains between HIR2-mCherry and YFP-REM1.2/YFP-REM1.3 proteins using Pearson’s correlation coefficient (PCC). Square regions of interest were analysed. As a control for random co-localization, the same regions where the YFP channel was rotated by 90° with respect to the mCherry channel were analysed (rot.). Scatter plots with indicated mean ± standard deviation. Letters indicate significant differences (p < 0.05, one-way ANOVA with post-hoc Tukey-Kramer multiple comparison test, n = 23, 28).

### HIR2 is mono S-acylated on either C6 or C7, which ensures its plasma membrane targeting

To determine how HIR proteins, that do not contain transmembrane domains, are targeted to and associate with the PM, we investigated the role of lipidations. Indeed, all four HIR isoforms were found to be N-terminally myristoylated on glycine 2 (Majeran et al., 2018). To verify that HIR2-mCherry fusion protein used in this study was indeed myristoylated, we immunopurified HIR2-mCherry expressed under the control of *HIR2* promoter in Arabidopsis seedlings and then analyzed HIR2-mCherry by LC-MS/MS (Supplemental Figure S3). We observed that the N-terminus of HIR2-mCherry fusion protein exhibited a truncation of the initiating methionine and myristoylation of the exposed glycine. Since no other form of the N-terminal tail could be detected, we assume that 100 % of the cellular pool of HIR2-mCherry is indeed myristoylated on G2. In addition to be myristoylated, HIR proteins were shown to be S-acylated, as determined by biotin switch assays (Hemsley et al., 2013). Predictions using GPS-Palm (Ning et al., 2021) suggested that in HIR2, C6 and C7 could be the palmitoylation sites (Figure 2U). An atlas of Arabidopsis protein S-acylation suggested that C58 in HIR2 was also S-acylated (Kumar et al., 2022). To study the role of myristoylation and S-acylation in HIR2 trafficking and nanodomain organization, amino acids G2, C6, C7 and C58 were mutated to alanine, individually or in combinations. WT and mutated HIR2 proteins fused to mCherry were transiently expressed in *N. benthamiana* leaf epidermal cells under the control of *HIR2* promoter (Figure 2A-J and Supplemental Figure S4). Confocal microscopy analysis revealed that the individual mutations G2A, C6A, C7A and C58A did not alter HIR2 PM localization (Figure 2B-E). By contrast, the C6A, C7A double mutation induced a strong intracellular retention of HIR2 protein (Figure 2F). A similar mis-localization was observed for HIR2^G2A, C6A, C7A^ or HIR2^C6A, C7A, C58A^ mutated proteins (Figure 2G, J). On the other hand, when C58A mutation was combined with either C6A or C7A mutations, HIR2^C6A, C58A^ and HIR2^C7A, C58A^ proteins were still localized at the PM (Figure 2H and I). Similar subcellular localizations were observed when WT and mutated HIR2 proteins fused to mCherry were stably expressed in *Arabidopsis thaliana* under the control of *HIR2* promoter (Figure 2K-T). Notably, confocal microscopy analysis performed on Arabidopsis root epidermal cells highlighted that the double mutation C6A, C7A induced an intracellular mis-localization of HIR2 (Figure 2P). Altogether these results suggest that: (i) myristoylation is not required for HIR2 PM targeting, (ii) C58 probably does not play a role in HIR2 trafficking, (iii) C6 and C7 together are essential to address HIR2 to the PM.

**Figure 2.**
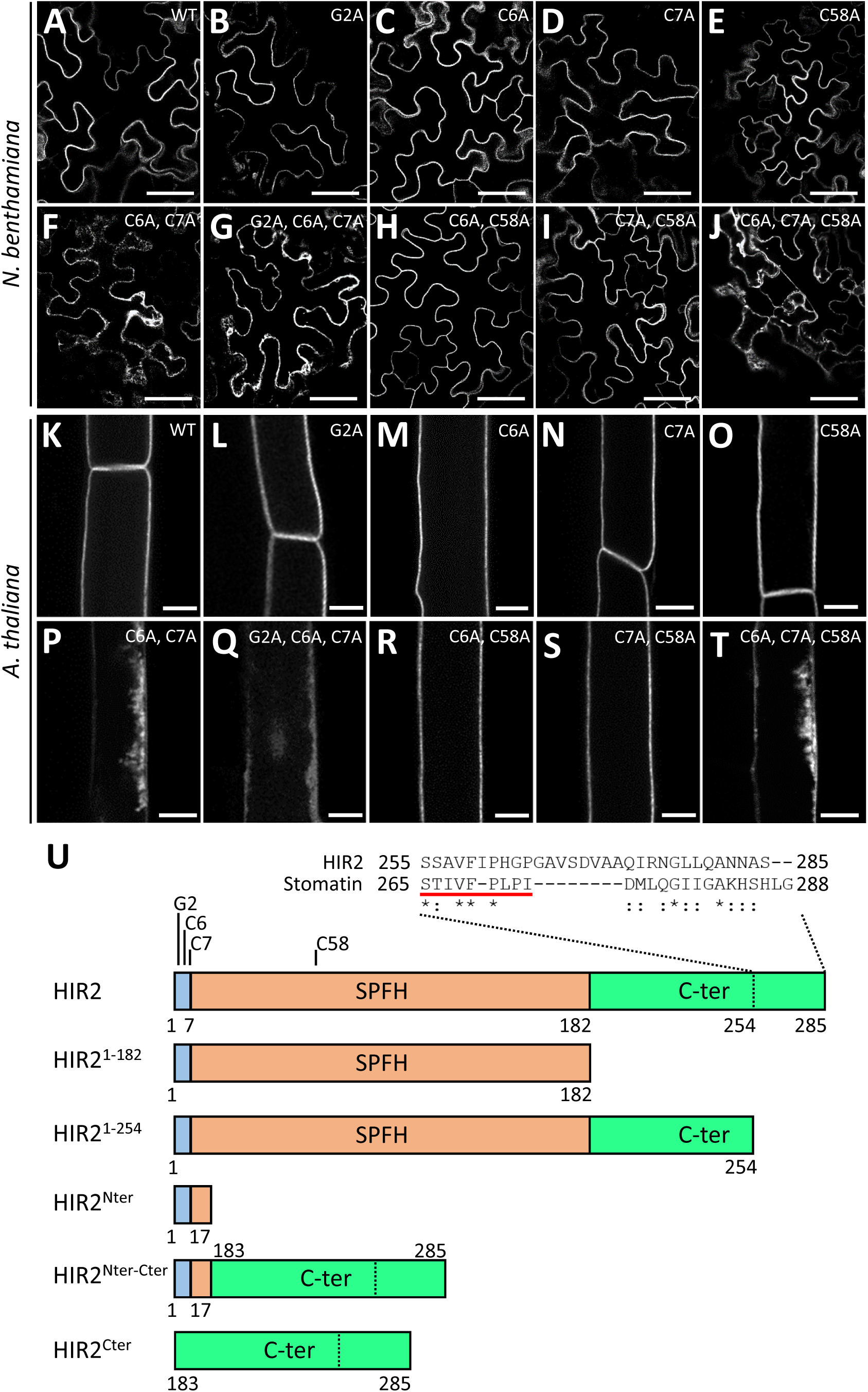
Simultaneous mutations of C6 and C7 induce HIR2 intracellular retention in plant cells. The subcellular localization of HIR2 point mutants fused to mCherry and expressed under the control of *HIR2* promoter in *N. benthamiana* leaf epidermal cells **(A-J)** and in the root epidermal cells of Arabidopsis transgenic lines **(K-T)** was analyzed by scanning confocal microscopy. **(A)** and **(K)**: HIR2-mCherry. **(B)** and **(L)**: HIR2^G2A^-mCherry. **(C)** and **(M)**: HIR2^C6A^-mCherry. **(D)** and **(N)**: HIR2^C7A^-mCherry. **(E)** and **(O)**: HIR2^C58A^-mCherry. **(F)** and **(P)**: HIR2^C6A, C7A^-mCherry. **(G)** and **(Q)**: HIR2^G2A, C6A, C7A^-mCherry. **(H)** and **(R)**: HIR2^C6A, C58A^ -mCherry. **(I)** and **(S)**: HIR2^C7A, C58A^-mCherry. **(J)** and **(T)**: HIR2^C6A, C7A, C58A^-mCherry. Secant views are shown. Arabidopsis plantlets were grown 11 days on MS/2 medium. Scale bars represent 50 µm and 10 µm in panels (A-J) and (K-T), respectively. **(U**) Schematic representation of HIR2 protein and the variants used in this work. HIR2 protein contains a N-terminal sequence composed of 7 amino acids, a SPFH domain (position 8 to 182) that was defined according to the Pfam website and a C-terminal domain (position 183 to 285). A sequence present in the very C-terminus of HIR2 displays homology to an oligomerization motif found in human Stomatin (underlined in red). Glycine 2 (G2) was shown to be myristoylated, cysteine 58 (C58) was proposed to be S-acylated and cysteines 6 and 7 (C6, C7) were predicted to be S-acylation sites according to GPS-Palm. Truncated variants of HIR2 and chimeric proteins are represented.

To analyze the role of S-acylation in HIR2 intracellular trafficking, we first used the S-acylation inhibitor 2-bromopalmitate (2-BP) that was successfully used to impair protein S-acylation in plant cells (Zhang et al., 2015). Although in mock-treated Arabidopsis seedlings HIR2-mCherry protein was only detected in the PM of root epidermal cells, 2-BP treatment induced a strong intracellular mis-localization of HIR2-mCherry (Figure 3A, right). In parallel, the subcellular localization of the PM protein GFP-LTi6b, which is not S-acylated, remained largely unchanged under 2-BP treatment, showing the specificity of 2-BP in this experiment. These results suggested that S-acylation is required for HIR2 trafficking to the PM in intact Arabidopsis root epidermal cells.

**Figure 3.**
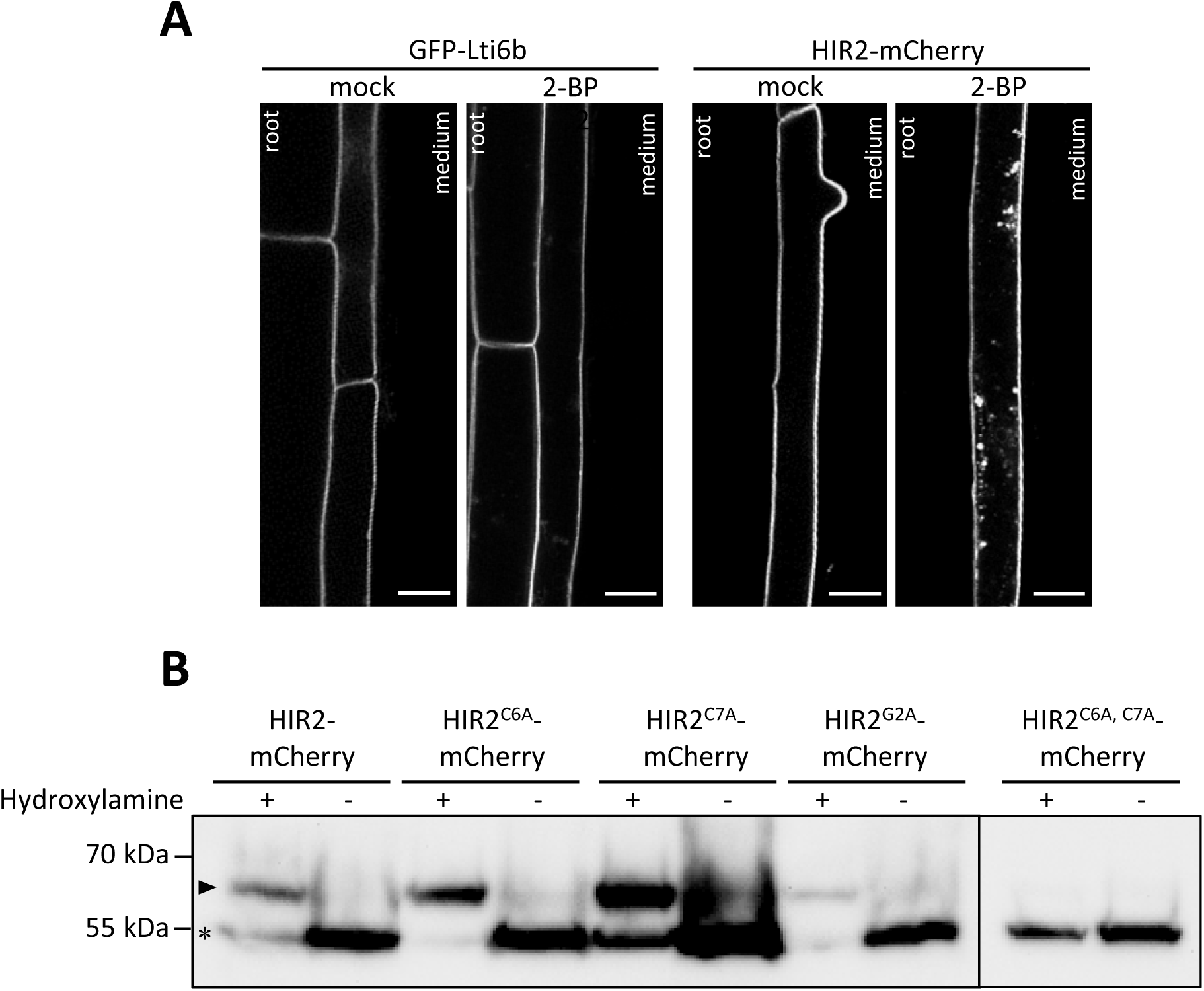
Mono S-acylation is implicated in HIR2 intracellular trafficking. **(A)** Inhibition of S-acylation using 2-bromopalmitate (2-BP) disturbs HIR2 subcellular localization. The subcellular localization of HIR2-mCherry and GFP-LTi6b (control) expressed under the control of *HIR2* and *35S* promoters, respectively, was analyzed by scanning confocal microscopy in the roots of 11-day-old Arabidopsis seedlings treated with 2-BP or mock treated (mock). Scale bars represent 20 µm. **(B)** HIR2 is mono S-acylated and simultaneous mutations of C6 and C7 abolish HIR2 S-acylation. The S-acylation of WT HIR2, HIR2^C6A^, HIR2^C7A^, HIR2^G2A^ and HIR2^C6A, C7A^ proteins fused to the mCherry and expressed under the control of *HIR2* promoter in Arabidopsis transgenic lines was assessed with a PEG-shift assay combined with anti-mCherry immunoblots. Following the cleavage of S-acyl groups with the thioester cleavage reagent hydroxylamine, a maleimide-functionalized polyethylene glycol (mPEG-Mal) was selectively coupled to free cysteines, which induced a mobility shift of 10 kDa for each S-acylation site. In the absence of hydroxylamine (-), no mass shift should be observed with a signal at ∼ 55 kDa, corresponding to the expected molecular weight of HIR2 and HIR2 mutated proteins fused to the mCherry. The mono S-acylation of WT HIR2, but also some HIR2 mutated proteins fused to the mCherry is indicated by the detection of a signal at ∼ 65 kDa in the presence of hydroxylamine (+). Non-S-acylated and mono S-acylated HIR2 and HIR2 mutated proteins fused to the mCherry are indicated by a star and an arrowhead, respectively. Arabidopsis seedlings were grown for 11 days on MS/2 medium.

Then, we analyzed the impact of C6A and C7A mutations, as well as G2A mutation, on HIR2 S-acylation by performing a PEG shift assay on Arabidopsis seedlings expressing WT HIR2 protein and HIR2 mutants fused to mCherry. The PEG shift assay allows to selectively replace S-fatty acids with maleimide-functionalized polyethylene glycol (mPEG-Mal) of defined mass, which induces a mobility shift of the protein proportional to the number of S-fatty acylated cysteines (Percher et al., 2017; Kumar et al., 2022). Here, we adapted the protocol described by Percher and co-workers (Percher et al., 2017). Briefly, following protein disulfide bond reduction and capping of non-modified cysteines with N-ethylmaleimide (NEM), cysteine-coupled fatty acid groups were specifically cleaved with hydroxylamine, followed by the coupling of 5 kDa mPEG-Mal. Note that the 5 kDa mPEG-Mal is known to induce in practice an apparent mass shift of 10 kDa (Percher et al., 2017). To control the efficiency of the capping of unmodified cysteines by NEM and avoid misinterpretations, an experiment was also performed in the absence of hydroxylamine (-Hyd). An anti-mCherry immunoblot revealed the presence of a single signal at the expected molecular weight for HIR2-mCherry (∼55 kDa) in - Hyd condition, whereas in the presence of hydroxylamine (+Hyd) a single mass shift of about 10 kDa was observed for HIR2-mCherry (∼65 kDa), suggesting that this protein is mono S-acylated (Figure 3B and Supplemental Figure S5). By contrast, no mass shift was observed for the HIR2^C6A, C7A^-mCherry mutated protein in the +Hyd condition, showing that C6A, C7A double mutation abolished HIR2 S-acylation. For both HIR2^C6A^-mCherry and HIR2^C7A^-mCherry, a mass shift of about 10 kDa was detected in +Hyd condition, suggesting that these two mutated proteins are still mono S-acylated (Figure 3B and Supplemental Figure S5). In +Hyd condition, a mass shift was also observed for HIR2^G2A^-mCherry, showing that G2A mutation does not interfere with HIR2 S-acylation. Altogether, these data strongly suggest that HIR2 is mono S-acylated on either C6 or C7 in Arabidopsis to ensure HIR2 PM targeting. Note that for HIR2, HIR2^C6A^, HIR2^C7A^ and HIR2^G2A^ fusion proteins, a signal was also detected in +Hyd at ∼55 kDa, which could indicate that not all the protein pools are S-acylated, but may also results from an incomplete binding of mPEG-Mal during the PEG shift assay. Intriguingly, the absence of mass shift for HIR2^C6A, C7A^ in +Hyd condition highly suggests, since C58 is not mutated in this protein, that C58 is not S-acylated in our experimental conditions.

### HIR2 plasma membrane targeting likely relies on mechanisms independent of the secretory pathway

To explore the localization of the intracellularly retained HIR2^C6A, C7A^ S-acylation mutant, we performed cell fractionation on *N. benthamiana* leaf tissues expressing HIR2^C6A, C7A^-mCherry. Surprisingly, similarly to WT HIR2, HIR2^C6A, C7A^ protein was exclusively detected in the microsomal fraction that corresponds to membranes and not in the cytosolic fraction (Supplemental Figure S6). Identical results were obtained for HIR2^G2A^, HIR2^C6A^ and HIR2^C7A^ mutants. In addition, HIR2^G2A, C6A, C7A^ triple mutant was still associated with membranes (Supplemental Figure S6). These data highly suggest that lipidations are not the sole way to anchor HIR2 into membranes and that HIR2 also interacts with membrane through other mechanisms.

Then, we aimed at better understanding the trafficking routes followed by HIR2 to reach the PM. We hypothesised that HIR2^C6A, C7A^ protein may be trapped in an intermediate compartment *en route* to the PM, possibly the ER. However, when HIR2^C6A, C7A^-mCherry was co-expressed with the pHluorine-HDEL ER marker in *N. benthamiana* leaf epidermal cells, no clear co-localization was observed (Supplemental Figure S7A-C), suggesting that mis-localized HIR2^C6A, C7A^ is not retained in the ER. In plants, the majority of PM localized proteins are synthesized in the ER and then traffic along the secretory pathway to ultimately reach the PM. These membrane proteins exit the ER through the action of the coat protein complex II (COPII) machinery that is recruited to the surface of the ER membrane to induce the formation of transport vesicles and ensure the delivery of cargo proteins to the Golgi apparatus (Chung et al., 2016). To investigate whether HIR2 PM delivery requires the COPII-mediated pathway, we used a dominant negative (DN) version of the COPII component SAR1 protein that is essential to initiate COPII vesicle budding (daSilva et al., 2004). As a positive control to monitor the effect of SAR1-DN, we used the Arabidopsis proton pump H^+^-ATPase2 (AHA2) that is mainly localized in the PM when expressed alone in *N. benthamiana* leaf epidermal cells (Supplemental Figure S7 D-E) and travels along the secretory pathway as other H^+^-ATPases (Lefebvre et al., 2004). Upon co-expression with SAR1-DN-YFP, AHA2-mCherry was fully retained in the ER, whereas HIR2-mCherry trafficking to the PM was only slightly disturbed, suggesting that HIR2 trafficking to the PM mainly occurs independently of COPII vesicles (Supplemental Figure S7 H-K).

### Single mutations of lipidation sites do not affect HIR2 plasma membrane nanodomain organization

Although G2A, C6A and C7A mutations did not affect HIR2 PM targeting, we wondered whether they could interfere with HIR2 nanodomain formation. To investigate HIR2 clustering, we used Total Internal Reflection Fluorescence (TIRF) microscopy that displays an enhanced signal to noise ratio due to shallow excitation of the cells compared to other microscopy techniques and constitutes one of the state of the art microscopy techniques to study nanodomain organization in the PM (Platre et al., 2019; Smokvarska et al., 2020). Importantly, we did not perform any imaging post-processing that can sometimes produce nanodomain-resembling artefacts (Jaillais and Ott, 2020). When expressed in *N. benthamiana* leaf epidermal cells or in Arabidopsis root epidermal cells, HIR2^G2A^, HIR2^C6A^ and HIR2^C7A^ mutated proteins fused to mCherry displayed a dotty like pattern in the PM, characteristic of nanodomain-organized proteins that was similar to the pattern of HIR2-mCherry (Figure 4A-D and 4H-K). To quantify the level of clustering of WT HIR2 and mutated HIR2 proteins in the PM, the coefficients of variance (CV) of pixel intensity that reflect here the heterogeneity of protein distribution in the PM, were calculated from TIRF acquisitions (Gronnier et al., 2017; Retzer et al., 2017; Danek et al., 2020). CV values calculated for WT HIR2, HIR2^G2A^, HIR2^C6A^ and HIR2^C7A^ proteins were not significantly different (Figure 4O). Therefore, we concluded that: (i) myristoylation is not involved in HIR2 nanodomain organization, (ii) single S-acylation mutants display a normal arrangement into nanodomains likely due to the interchangeability of S-acylation sites, C6 or C7. Although the PEG shift assay highly suggested that C58 is not S-acylated in our experimental conditions, we also tested the clustering of HIR2^C58^, HIR2^C6A, C58A^ and HIR2^C7A, C58A^ mutated proteins. TIRF analysis revealed that all these mutated proteins displayed a normal arrangement into PM nanodomains (Figure 4E-G, 4L-N and 4O), showing that C58 residue has no role in HIR2 nano-organization, nor in its PM targeting as mentioned above.

**Figure 4.**
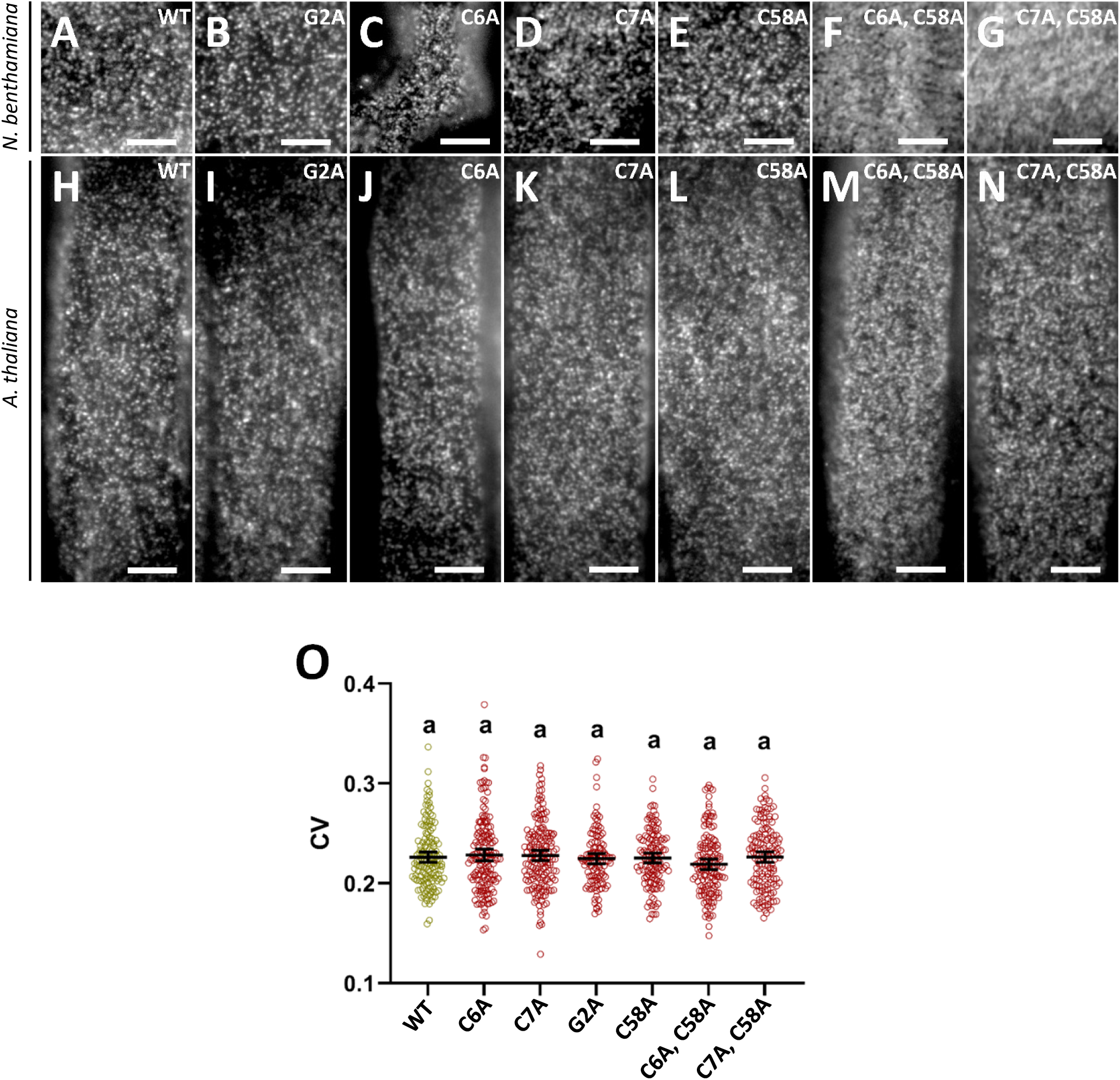
HIR2 lipidation mutants that are targeted to the plasma membrane still organize in nanodomains. Total Internal Reflection Fluorescence (TIRF) microscopy analysis was performed on the PM of *Nicotiana benthamiana* leaf epidermal cells **(A-G)** or *Arabidopsis thaliana* root epidermal cells **(H-O)** expressing HIR2 point mutants fused to mCherry and expressed under the control of *HIR2* promoter. Note that the arrangement in nanodomains was only investigated for HIR2 mutated proteins that are still targeted/localized to the PM. **(A)** and **(H)**: HIR2-mCherry. **(B)** and **(I)**: HIR2^G2A^-mCherry. **(C)** and **(J)**: HIR2^C6A^-mCherry. **(D)** and **(K)**: HIR2^C7A^-mCherry. **(E)** and **(L)**: HIR2^C58A^-mCherry. **(F)** and **(M)**: HIR2^C6A, C58A^-mCherry. **(G)** and **(N)**: HIR2^C7A, C58A^-mCherry. **(O)** Coefficients of variance (CV) of fluorescence intensity were calculated from TIRF experiments performed as in (H) to (N) on Arabidopsis root epidermal cells. Error bars correspond to a confidence interval at 95%. Letters indicate significant differences (p-value < 0.05, one-way ANOVA with post-hoc Tukey multiple comparison test, n>25). Arabidopsis plantlets were grown 7 days on MS/2 medium. Scale bars represent 5 µm.

### Sterols and very long chain fatty acids influence HIR2 organization in the plasma membrane

To investigate the importance of PM lipid composition on HIR2 nanodomain organization, we focused our attention on: (i) sterols that play a crucial role in the structuration of nanodomains, (ii) VLCFA, since PS and sphingolipids, which both contain VLCFA, are closely linked to the organization of nanodomains (Grosjean et al., 2015; Gronnier et al., 2017; Platre et al., 2019). First, we used the sterol synthesis inhibitor fenpropimorph (FEN) that alters the PM sterol composition but not the total amount of sterols (Gronnier et al., 2017). Following FEN treatment, HIR2-mCherry clustering was decreased in the PM of Arabidopsis root epidermal cells compared to non-treated or mock treated plants, as revealed by TIRF microscopy (Figure 5A-C). The negative effect of FEN on HIR2 clustering was illustrated by a reduction of the CV value (Figure 5E). A similar effect of FEN was observed on the nanodomain organization of GFP-StREM1.3, as previously shown (Supplemental Figure S8) (Gronnier et al., 2017). Second, to alter the composition of VLCFA incorporated in the pool of sphingolipids and phosphatidylserine in the PM, we used metazachlor (MTZ) that reduces the length of the chains of VLCFA by targeting the 3-ketoacyl CoA synthase enzymes (Wattelet-Boyer et al., 2016). MTZ treatment dramatically changed the localization of HIR2-mCherry that displayed a homogeneous signal in the PM of Arabidopsis root epidermal cells (Figure 5D) different from the controls where HIR2-mCherry was clustered (Figure 5A-B). This was accompanied by a strong reduction of the CV value upon MTZ treatment (Figure 5E). To ensure that MTZ treatment did not induce a general disorganization of the PM, we showed that the nanodomain organization of GFP-StREM1.3 expressed in Arabidopsis root epidermal cells was not affected by MTZ (Supplemental Figure S9). Altogether, these results showed that both sterols and VLCFA are important for the nanodomain organization of HIR2 in the PM.

**Figure 5.**
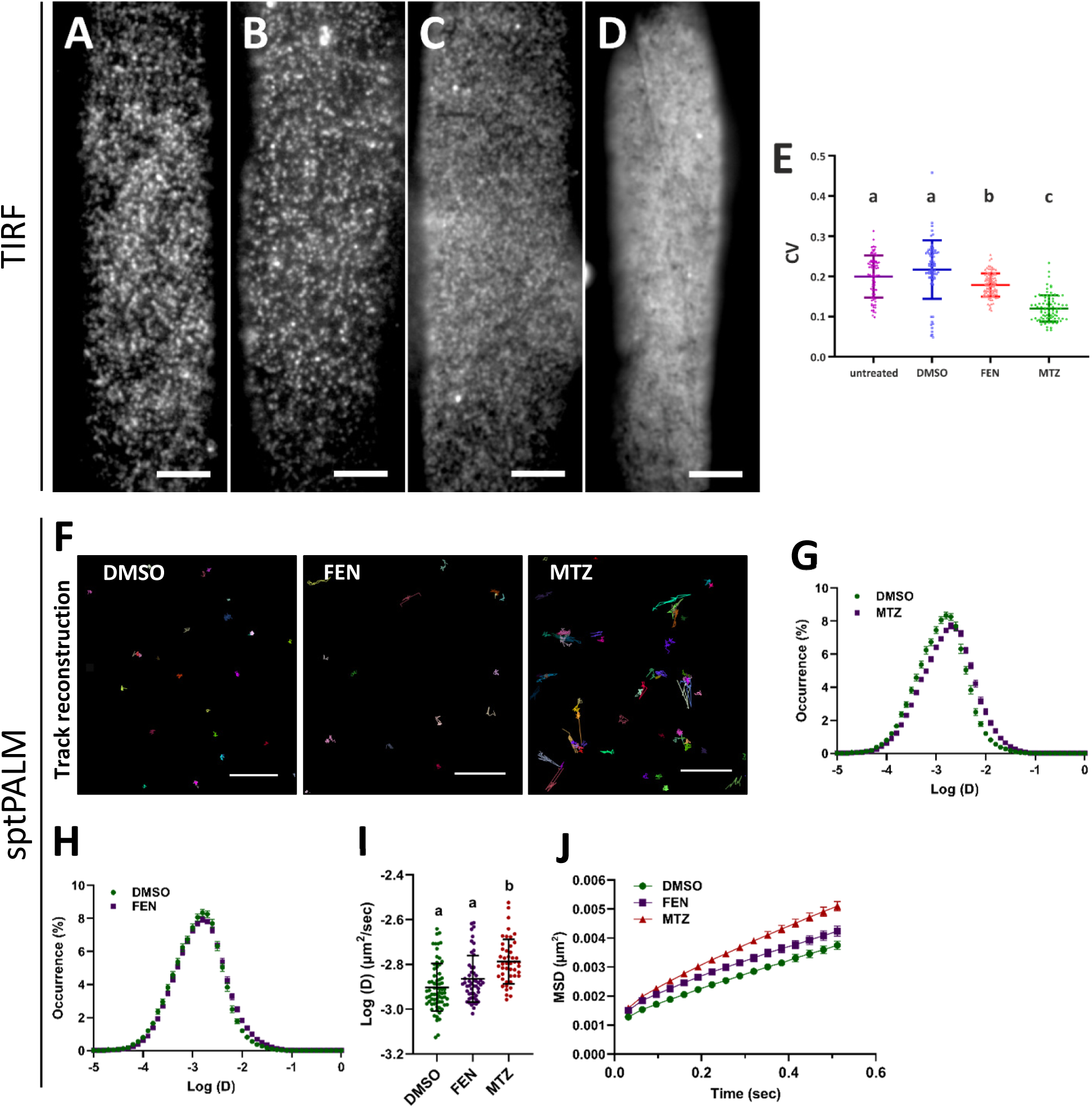
Sterols and very long chain fatty acids (VLCFA) are involved in the organization of HIR2 in the plasma membrane. Total Internal Reflection Fluorescence (TIRF) microscopy analysis on the PM of root epidermal cells from pHIR2::HIR2-mCherry transgenic Arabidopsis seedlings, non-treated **(A)**, mock-treated with DMSO **(B)**, treated with fenpropimorph (FEN) **(C)** or metazachlor (MTZ) **(D)**. Scale bars represent 5 µm. (**E)** Coefficient of variance (CV) of fluorescence intensity determined from TIRF images of root epidermal cells expressing HIR2-mCherry and treated as in (A-D). Scatter plots with indicated mean ± standard deviation. Letters indicate significant differences (p < 0.05, one-way ANOVA with post-hoc Tukey-Kramer multiple comparison test, n = 70 – 100). (**F-J)**: Single-particle-tracking photoactivated localization microscopy (sptPALM) was performed on the PM of root epidermal cells from pHIR2::HIR2-mEOS2 mock-treated Arabidopsis seedlings (DMSO) or seedlings treated with FEN or MTZ. **(F)** For each treatment, image reconstructions of several tens of single HIR2-mEOS2 molecule trajectories are indicated by different colours. Scale bars represent 10 µm. The distribution of HIR2-mEOS2 molecule instantaneous diffusion coefficient (D) (expressed in log (µm^2^/sec)) for the different treatments is presented in **(G)** and **(H)**. **(I)** Quantification of the diffusion of HIR2-mEOS2 under the different treatments. Letters indicate significant differences (p < 0.05, one-way ANOVA with post-hoc Tukey multiple comparison test, n = 50 – 60). **(J)** Mean square displacement (MSD) curves of HIR2-mEOS2 molecules under the different treatments.

To analyze deeper the effect of the disruption of the sterol and VLCFA composition on HIR2 dynamics in the PM, we performed single-particle-tracking photoactivated localization microscopy (sptPALM) analysis, using an Arabidopsis transgenic line expressing HIR2 fused to the mEOS2 photoconvertible fluorescent protein under the control of *HIR2* promoter. sptPALM is a super-resolution technique allowing to visualize and track single molecules along time in living cells and measure different kinetic and organizational parameters including instantaneous diffusion (D) and mean square displacement (MSD) (Hosy et al., 2015; Gronnier et al., 2017; Martiniere et al., 2019). Upon mEOS2 stochastic photoactivation, sub diffraction spots appear with a blinking behaviour, as expected from single molecules (Supplemental Video 1). Typically, several thousands of single HIR2-mEOS2 molecules were imaged from a single time series. The localization of molecules along time was retrieved and tracks were reconstructed with a spatial resolution of ∼30 nm and a temporal resolution of 32 ms. Then, tracks were used to infer molecule instantaneous diffusion (Bayle et al., 2021) (Figure 5F-J). MTZ treatment increased the diffusion of HIR2-mEOS2 molecules in the PM of root epidermal cells (D = 0.003 ± 0.0001 µm^2^ s^-1^) compared to mock-treated conditions (D =0.002 ± 0.0006 µm^2^ s^-1^) (Figure 5F, G, I). In addition, the MSD of HIR2-mEOS2 was faster in presence of MTZ, showing an increased mobility of HIR2-mEOS2 molecules in the PM (Figure 5J). By contrast, FEN treatment induced a very slight but not significant increase of D and MSD parameters, showing that HIR2-mEOS2 diffusion was not affected by FEN (Figure 5F, H, I, J).

### Oligomerization of HIR2 through its C-terminal domain is required for HIR2 nano-organization in the plasma membrane

Since HIR2 protein is known to homo-oligomerize (Qi et al., 2011), we analyzed the effect of the co-expression of WT HIR2 protein with the HIR2^C6A, C7A^ mutant that is retained intracellularly. Interestingly, upon co-expression with HIR2-mEGFP, HIR2^C6A, C7A^-mCherry was relocalized from intracellular compartments to the PM as revealed by confocal microscopy (Supplemental Figure S10A). In addition, HIR2-mEGFP and HIR2^C6A, C7A^ -mCherry proteins were strongly co-localized in clusters at the PM (Supplemental Figure S10B-C). Therefore, the expression of WT HIR2 protein can drive non-acylated HIR2 mutated proteins to PM nanodomains.

We sought to identify the domains involved in HIR2 homo-oligomerization, this latter being probably important for HIR2 function. Interestingly, HIR2 protein carries in its very C-terminal part a sequence that displays a partial identity with an oligomerization motif present in the human SPFH domain-containing protein Stomatin (Figure 2U) (Umlauf et al., 2006). We deleted this motif to generate the truncated HIR2^1-254^ protein (amino acids 1-254) and also created the HIR2^1-182^ truncated protein in which the whole C-terminal part was deleted to conserve the N-terminal sequence and the SPFH domain of HIR2 (amino acids 1-182) (Figure 2U). When expressed in *N. benthamiana* leaf epidermal cells and in Arabidopsis root epidermal cells under the control of *HIR2* promoter, both HIR2^1-254^ and HIR2^1-182^ proteins fused to fluorophores were solely observed in the PM, as WT HIR2, showing that the C-terminal part of HIR2 is not required for PM targeting (Figure 6A-I). To analyze the effect of HIR2 C-terminal truncations on HIR2 oligomerization, we performed co-immunopurifications (co-IP) between HIR2, HIR2^1-254^, HIR2^1-182^ proteins fused to mEGFP and HIR2-Cherry expressed in *N. benthamiana* leaf epidermal cells. Anti-mCherry immunoblots revealed that HIR2-mCherry protein is co-purified with HIR2-mEGFP but not with HIR2^1-182^-mEGFP (Figure 6J and Supplemental Figure S11), showing that this truncated protein lost its capacity to interact with other HIR2 molecules. In addition, a faint mCherry signal was detected in HIR2^1-254^-mEGFP IP fraction, indicating a weaker interaction between HIR2-mCherry and HIR2^1-254^-mEGFP, compared to HIR2-mEGFP. These results highlighted that the C-terminal part of HIR2 is required for HIR2-HIR2 protein interaction. In accordance with these interaction tests, co-expression analysis in *N. benthamiana* leaf epidermal cells showed that, in contrast to HIR2-mEGFP, HIR2^1-182^-mEGFP protein is not able to induce a relocalization of HIR2^C6A, C7A^-mCherry to the PM (Supplemental Figure S12A-F), whereas a partial targeting of this protein to the PM occurs upon co-expression with HIR2^1-254^-mEGFP (Supplemental Figure S12G-I).

**Figure 6.**
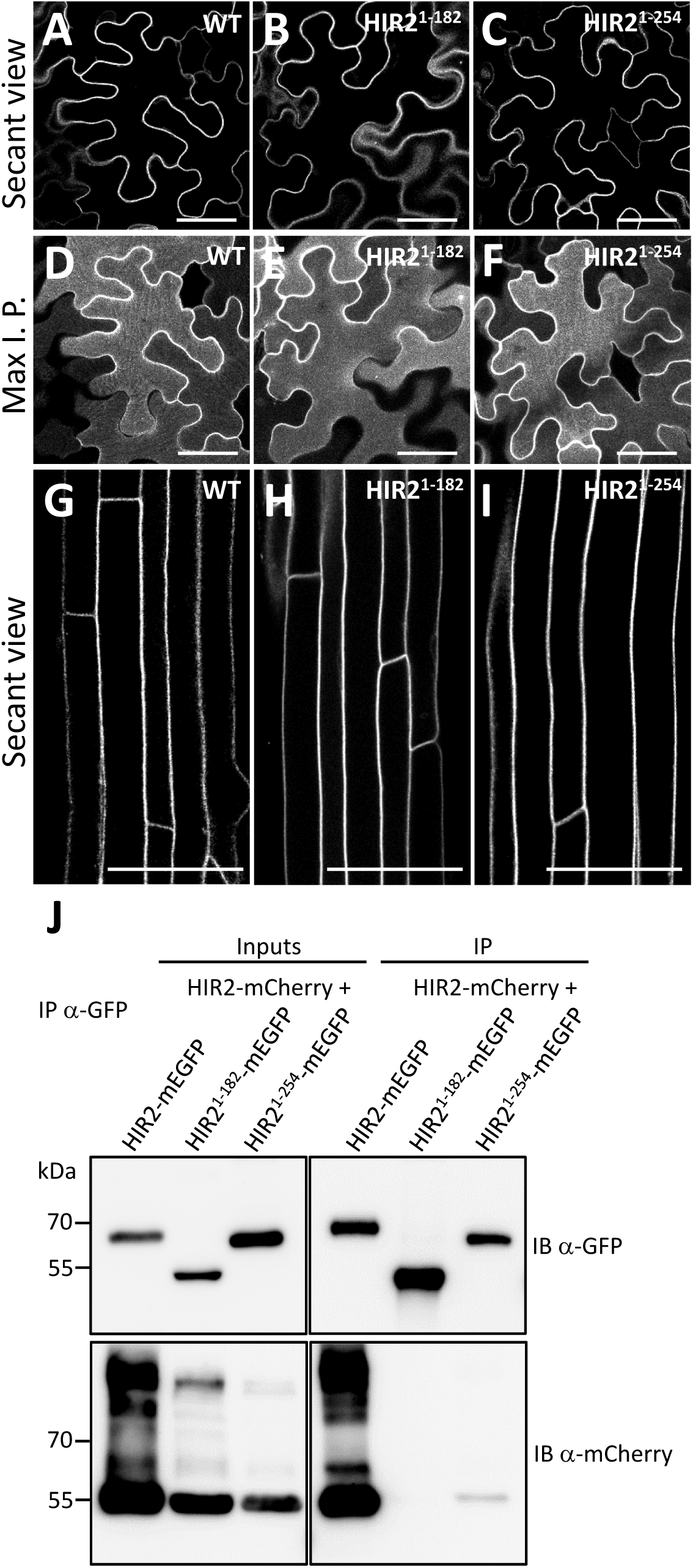
The C-terminal part of HIR2 is required for oligomerization but not for plasma membrane targeting. The subcellular localization of HIR2-mEGFP **(A, D)**, HIR2^1-182^-mEGFP **(B, E)** and HIR2^1-254^-mEGFP **(C, F)** transiently expressed under the control of *HIR2* promoter in *N. benthamiana* leaf epidermal cells was analyzed by scanning confocal microscopy. HIR2^1-182^ and HIR2^1-254^ are truncated proteins corresponding to the first 182 and 254 amino acids of HIR2, respectively. (A-C): Secant views. (D-F): Maximum intensity projections constructed from Z-stack of 25 focal planes spanning from the epidermis surface to the positions shown in (A-C). **(G-I)**: Secant views of root epidermal cells from Arabidopsis transgenic lines expressing HIR2-mCherry **(G)**, HIR2^1-182^-mCherry **(H)** and HIR2^1-254^-mEGFP **(I)** under the control of *HIR2* promoter. Scanning confocal microscopy, scale bars represent 50 µm. Arabidopsis plantlets were grown 11 days on MS/2 medium. **(J)** Deletions in the C-terminal part of HIR2 disturb HIR2 oligomerization as revealed by co-immunopurifications. Immunopurifications (IP) using an anti-GFP antibody were performed on solubilized protein extracts from *N. benthamiana* leaves transiently co-expressing HIR2-mCherry with HIR2-mEGFP, HIR2^1-182^-mEGFP or HIR2^1-254^-mEGFP. Inputs and IP fractions were subjected to immunoblots (IB) with anti-GFP (top) and anti-mCherry antibodies (bottom).

To investigate the importance of oligomerization in HIR2 nanodomain organization, we performed TIRF microscopy on the PM of *N. benthamiana* leaf epidermal cells and Arabidopsis root epidermal cells expressing HIR2, HIR2^1-182^ and HIR2^1-254^ proteins fused to fluorophores. Contrary to WT HIR2 that arranged in nanodomains (Figure 7A, D), HIR2^1-182^ displayed a diffuse localization in the PM (Figure 7B, E) illustrated by a strong reduction of its CV value (Figure 7G). An intermediate situation was observed for HIR2^1-254^ that partially arranges in nanoclusters (Figure 7C, F) as attested by a CV value between the ones of WT HIR2 and HIR2^1-182^ (Figure 7G). Then, we wondered if this massive change in the localization of HIR2^1-182^ protein could be associated with a change in its dynamics in the PM. Therefore, HIR2 and HIR2^1-182^ fused to mEOS2 were expressed in *N. benthamiana* leaf epidermal cells under the control of *HIR2* promoter, followed by sptPALM analysis. The localization and displacement of single molecules of WT HIR2 and HIR2^1-182^ were reconstructed and used to assess their instantaneous diffusion (Figure 7H, J-L). The diffusion of HIR2^1-182^-mEOS2 protein (D =0.042 ± 0.004 µm^2^ s^-1^) was much higher than the one of HIR2-mEOS2 (D =0.0016 ± 0.0001 µm^2^ s^-1^) (Figure 7H, J, K). MSD values were higher for HIR2^1-182^-mEOS2 compared to HIR2-mEOS2 showing that HIR2^1-182^ protein is more mobile than HIR2 in the PM (Figure 7L). In addition, clustering analysis on PALM images using Vonoroï tessellation showed that the occurrence of molecules with high local density was higher for HIR2-mEOS2 compared to HIR2^1-182^-mEOS2 (Figure 7I, M). These combined approaches of TIRF and sptPALM strongly suggest that interfering with HIR2 oligomerization affects HIR2 nanoclustering and diffusion in the PM.

**Figure 7.**
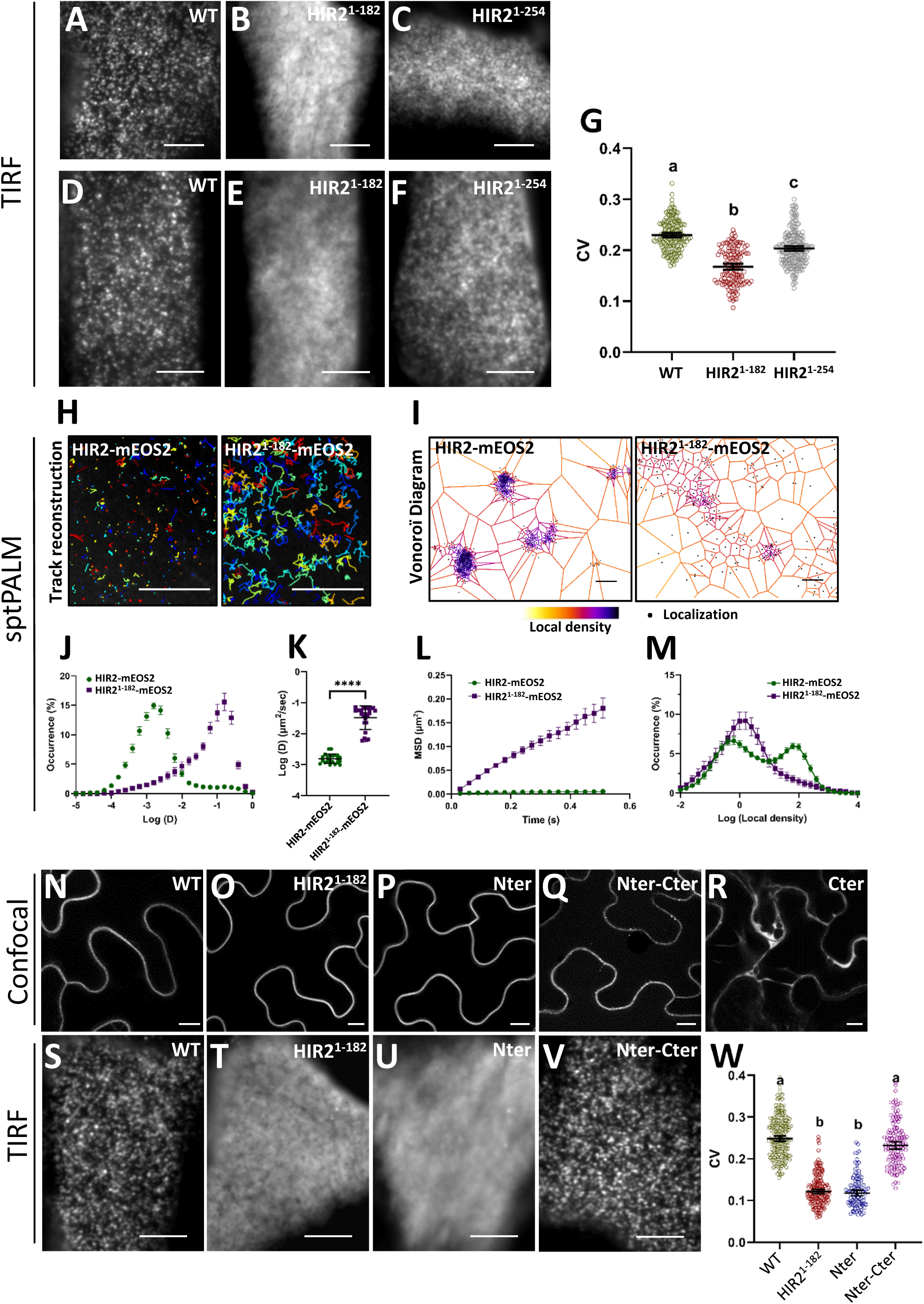
Oligomerization of HIR2 is required for its nano-organization in the plasma membrane. The organization in the MP of HIR2-mEGFP **(A, D)**, HIR2^1-182^-mEGFP **(B, E)** and HIR2^1-254^-mEGFP **(C, F)** proteins expressed under the control of *HIR2* promoter was analysed by TIRF microscopy. **(A-C)**: *N. benthamiana* leaf epidermal cells. **(D-F)**: Arabidopsis root epidermal cells. Arabidopsis seedlings were grown for 7 days on MS/2 medium. **(G)** Coefficients of variance (CV) of fluorescence intensity were calculated from TIRF experiments performed as in (D) to (F) on Arabidopsis root epidermal cells. Error bars correspond to a confidence interval of 95%. Letters indicate significant differences (p < 0.05, one-way ANOVA with post-hoc Tukey multiple comparison test, n>20). **(H-M)** Single-particle-tracking photoactivated localization microscopy (sptPALM) was performed on the PM of *N. benthamiana* leaf epidermal cells transiently expressing HIR2-mEOS2 and HIR2^1-182^-mEOS2 to determine the trajectories of each single protein. **(H)** Image reconstruction of several tens of single HIR2-mEOS2 and HIR2^1-182^-mEOS2 molecule trajectories indicated by different colors. **(I)** Vonoroï tessellation of HIR2-mEOS2 and HIR2^1-182^-mEOS2 molecule localization map from the same cells as in (H). The color code represents the local density of each molecule indicated by a black point. **(J)** Distribution of molecule instantaneous diffusion coefficient (D) (expressed in log (µm^2^/sec)) for HIR2-mEOS2 and HIR2^1-182^-mEOS2. **(K)** Quantification of the diffusion of HIR2-mEOS2 and HIR2^1-182^-mEOS2 molecules. **(L)** Mean square displacement (MSD) curves of HIR2-mEOS2 and HIR2^1-182^-mEOS2 molecules over the time. **(M)** Distribution of molecule local density for HIR2-mEOS2 and HIR2^1-182^-mEOS2. The subcellular localization of HIR2-mCherry **(N)**, HIR2^1-182^-mCherry **(O)**, HIR2^Nter^-mEGFP **(P)**, HIR2^Nter-Cter^-mCherry **(Q)** and HIR2^Cter^-mCherry **(R)** transiently expressed under the control of *HIR2* promoter in *N. benthamiana* leaf epidermal cells was analyzed by scanning confocal microscopy (secant views). HIR2^Nter^ and HIR2^Cter^ corresponds to the first 17 amino acids and the last 103 amino acids of HIR2, respectively, HIR2^Nter-Cter^ is a fusion of the first 17 amino acids of HIR2 to the last 103 amino acids. **(S-V)** TIRF microscopy analysis on the PM of *N. benthamiana* leaf epidermal cells expressing the same proteins than in (N) to (Q). **(W)** Coefficients of variance (CV) of fluorescence intensity were calculated from TIRF experiments performed as in (S-V). In this figure, error bars correspond to a confidence interval at 95% (G, W) and to standard deviation (K). Letters indicate significant differences (p-value < 0.05, one-way ANOVA with post-hoc Tukey multiple comparison test, n>20). In (K), **** represents p< 0.0001 in a t-test. Scale bars represent 5 µm (A-F ; S-V), 10 µm (H, N-R) and 0.102 µm (I).

To investigate deeper the role of the C-terminal part of HIR2 in nanodomain organization, we generated additional truncated and chimeric HIR2 proteins: (1) HIR2^Nter^ that corresponds to the 17 first amino acids of HIR2 including the S-acylation sites, (2) HIR2^Nter-Cter^, a fusion between the 17 first amino acids and the C-terminal part of HIR2 (amino acids 183-285), (3) HIR2^Cter^ that corresponds to the C-terminal part of HIR2 (Figure 2U). When expressed in *N. benthamiana* leaf epidermal cells, HIR2^Nter^-mEGFP solely displayed a PM localization as revealed by confocal microscopy, showing that a minimal sequence of HIR2 containing S-acylation sites is sufficient to ensure PM targeting (Figure 7P). HIR2^Nter-Cter^-mCherry was targeted to the PM, although it was also detected in rare intracellular vesicles (Figure 7Q). By contrast, HIR2^Cter^-mCherry displayed a cytosolic localization (Figure 7R). TIRF microscopy revealed that similarly to HIR2^1-182^, HIR2^Nter^ fused to mEGFP displayed a diffuse localization in the PM, which was rather expected since this truncated protein does not carry the C-terminal part of HIR2 required for oligomerization and clustering (Figure 7T, U). Thus, HIR2^1-182^ and HIR2^Nter^ proteins displayed similar CV values that were much lower than the one measured for WT HIR2 (Figure 7W). Interestingly, HIR2^Nter-Cter^-mCherry protein was detected in PM clusters and the corresponding CV value was not different from the one of WT HIR2 (Figure 7V, W). This result showed that the addition of the C-terminal part of HIR2 to a PM targeted fluorescent protein is sufficient to induce its arrangement in nanodomains. Based on this set of data, we propose that HIR2 oligomerization through its C-terminal part is essential for HIR2 organization in PM nanodomains.

### HIR2 nanodomain organization is required for HIR2-mediated ROS burst in response to flg22

In various organisms, a rapid burst of ROS is a conserved process during the immune response. Interestingly, the overexpression of pepper CaHIR1 protein in Arabidopsis can induce the accumulation of H_2_O_2_ in plant tissues (Jung and Hwang, 2007) and, recently, the ROS burst triggered by flg22 was shown to be attenuated in Arabidopsis *hir2* mutant (Liu et al., 2024). To investigate whether HIR2 nanodomain organization is necessary for HIR2 functioning, we compared the ability of WT HIR2 and HIR2^1-182^, which is still present in the PM but does not organize in nanodomains, to be implicated in the ROS burst in response to flg22. To do so, we quantified the apoplastic ROS production in tobacco leaf discs expressing either HIR2-mCherry, HIR2^1-182^-mCherry or mCherry (negative control) under the control of *HIR2* promoter, treated or not with flg22, using a luminol/peroxidase-based method. ROS accumulation was measured by following the emitted luminescence that was normalized, for each leaf disc, to mCherry fluorescence intensity, i.e. to the expression of the proteins of interest. Upon flg22 treatment, leaf discs expressing HIR2-mCherry displayed a rapid and higher ROS accumulation compared to leaf discs expressing mCherry (Figure 8A and B). Interestingly, in response to flg22 stimulus, the production of ROS in leaf discs expressing HIR2^1-182^-mCherry was much weaker than for HIR2-mCherry and not significantly different from the one observed in the negative control mCherry (Figure 8B). We concluded that HIR2 ectopic expression boosted apoplastic ROS accumulation in plant cells in response to flg22 and that this effect was cancelled when HIR2 lost the ability to oligomerize and to organize in PM clusters (HIR2^1-182^). Therefore, HIR2 nanodomain organization is a prerequisite for proper HIR2 functions in the cell.

**Figure 8.**
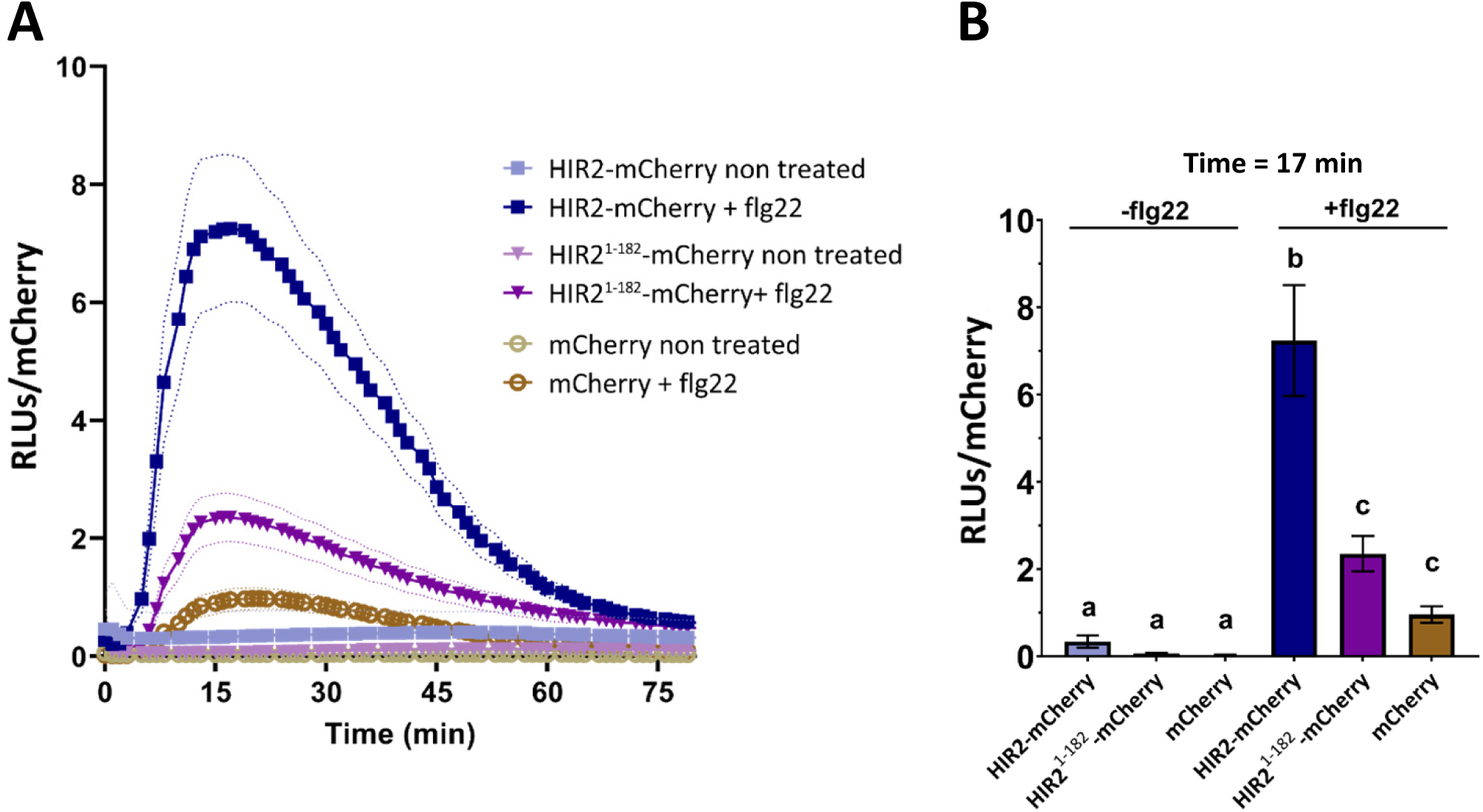
HIR2-stimulated oxidative burst in response to flg22 required HIR2 nanodomain organization. **(A-B)** The production of apoplastic ROS was measured with a luminol-based assay in *Nicotiana benthamiana* leaf discs expressing the following proteins: HIR2-mCherry, HIR2^1-182^-mCherry and mCherry (negative control). **(A)** Leaf discs were treated or not with 100 nM flg22 and ROS accumulation was measured by following the emitted luminescence, every 1 min for 80 min. The emitted luminescence was normalized to the quantity of HIR2-mCherry, HIR2^1-182^-mCherry and mCherry, estimated via mCherry fluorescence, and expressed in relative light units (RLUs). Curves show mean values with discontinued lines representing SE. A representative experiment among three independent biological experiments is shown. **(B)** The average ROS accumulation at 17 min (maximum of the peak) was measured for three independent biological experiments performed as in (A). Error bars represent SE. Letters indicate statistically significant differences between the different proteins that were expressed (ANOVA, multiple comparisons post-hoc Tukey’s method at P-value<0.05). For each protein, 12 and 22 leaf discs for non-treated and flg22-treated conditions were analyzed, respectively.

## DISCUSSION

### Mono S-acylation of HIR2 is required for its targeting to the PM independently of the conventional secretory pathway

In plants, the PM is sub-compartmentalized in a multitude of diverse nanodomains corresponding to the functional assemblies of specific proteins and lipids that act as signaling/regulation platforms (Jarsch et al., 2014; Smokvarska et al., 2020; Hdedeh et al., 2025). How these nanodomains are formed and maintained in the PM remains largely unknown. Here, we investigated the mechanisms governing the trafficking and the organization into PM nanodomains of the plant-specific SPFH domain-containing protein HIR2.

We analyzed the role of two types of post-translational modifications, myristoylation and S-acylation, in HIR2 intracellular dynamics and nanodomain organization in Arabidopsis. Pharmacological treatment interfering with HIR2 S-acylation led to mis-localization of HIR2-mCherry protein, expressed under the control of *HIR2* endogenous promoter, in Arabidopsis root epidermal cells (Figure 3). In addition, we showed that HIR2^C6A, C7A^ mutated protein is no more S-acylated in Arabidopsis transgenic lines (Figure 3), which triggers its retention inside the cell (Figure 2). Similar results were previously obtained by transiently overexpressing HIR2^C6S, C7S^-GFP in Arabidopsis protoplasts (Liu et al., 2021). However, it is important to mention that caution should be taken with protoplasts when studying PM nanodomain-organized proteins, because the cell wall plays an important role in the dynamics of such proteins, including HIR (Hosy et al., 2015; McKenna et al., 2019; Danek et al., 2020). Interestingly, we revealed using a PEG shift assay that HIR2 is mono S-acylated on either C6 or C7 (Figure 3). This interchangeability in HIR2 S-acylation sites probably explain why HIR2^C6A^ and HIR2^C7A^ mutated proteins display normal PM targeting and nanodomain organization (Figure 2 and 3). In addition, we showed that the very N-terminal part of HIR2 that carries the S-acylation sites is sufficient to drive a GFP protein to the PM (Figure 7), which strengthens the role of S-acylation in the trafficking of HIR2. Why HIR2 is mono S-acylated whereas both C6 and C7 can be S-acylation sites remains an open question. Interestingly, a site-specific analysis of neuronal protein S-acylation in mouse revealed that S-acylation sites on transmembrane proteins were enriched for di-cysteine motifs (CC), each cysteine could be S-acylated (Collins et al., 2017). We also analyzed a putative implication of C58 in HIR2 dynamics since this cysteine residue was proposed to be S-acylated (Kumar et al., 2022). However, in our experimental conditions, C58 likely did not undergo S-acylation (Figure 3) and had neither role in HIR2 PM targeting, nor in its nanodomain organization (Figures 2 and 4). Intriguingly, HIR2^C6A, C7A^ mutated protein still associated with intracellular membranes (Supplemental Figure S6), suggesting that lipidations are not the sole way to anchor HIR2 in membranes and that HIR2 may contain specific lipid interaction motifs. Indeed, some animal SPFH domain-containing proteins bind to sterols (Huber et al., 2006) and HIR2 was identified among Arabidopsis proteins interacting with the fungal sterol ergosterol (Khoza et al., 2019). We noticed that WT HIR2 protein could induce the relocalization of HIR2^C6A, C7A^ to PM nanodomains via oligomerization (Supplemental Figure S10), showing that in these experimental settings not all the HIR2 subunits need to be S-acylated among a complex to be targeted to the PM. Whether HIR2 S-acylation is systematic in plant cells or concerns a part of HIR2 subunits remains at this stage an open question. In our PEG-shift assay, we observed that a part of WT HIR2 protein did not appear to be S-acylated (Figure 3), but, as previously mentioned, we cannot exclude that it may result from an incomplete binding of mPEG-Mal during the PEG shift procedure.

HIR2 was demonstrated to be myristoylated on G2 (Majeran et al., 2018), a result we confirmed here by mass spectrometry using HIR2-mCherry fusion protein (Supplemental Figure S3). Some proteins bear both myristoylation and proximal S-acylation in their N-terminal part and this combination of lipidations is required for their PM targeting as exemplified by the plant calcineurin B–like protein 1 (CBL1) (Batistic et al., 2008). Indeed, both CBL1 myristoylation and S-acylation mutants are not properly addressed to the PM and myristoylation and S-acylation were proposed to be responsible for CBL1 targeting to the ER and for CBL1 trafficking from the ER to the PM, respectively. Interestingly, myristoylation can be a prerequisite for S-acylation, as shown for the plant GTPase Ara6 (Ueda et al., 2001). Here, we observed that HIR2^G2A^ mutated protein was properly targeted to the PM (Figure 2) and displayed a similar S-acylation profile compared to WT HIR2 (Figure 3), ruling out the possibility that HIR2 myristoylation influences its S-acylation. In addition, the normal clustering pattern of HIR2^G2A^ mutated protein in the PM excluded a possible role of this lipidation in HIR2 nanodomain organization (Figure 4). The absence of a role of myristoylation on HIR2 localization raises the question of the function of this modification. Interestingly, apart from membrane association, myristoylation is also involved in protein-protein interaction and protein activation (Wright et al., 2011; Zhu et al., 2019), giving some clues to explore the cellular function of HIR2 myristoylation in the future.

Investigating HIR2 intracellular trafficking, we highlighted that HIR2 PM targeting likely relies on mechanisms independent of the conventional secretory pathway involving COPII machinery (Supplemental Figure S7), similarly to what was shown for the nanodomain organized protein StREM1.3 (Gronnier et al., 2017). However, StREM1.3 was proposed to be targeted to the PM from the cytosol whereas our results suggest that HIR2 traffics through an intracellular membrane compartment, identity of which remains to be determined, before reaching the PM. Indeed, we believe that the membranes where HIR2^C6A, 67A^ protein is trapped and that largely differs from the ER is probably on the natural pathway taken by HIR2 to reach the PM since WT HIR2 can encounter HIR2^C6A, 67A^ and relocalize it to the PM (Supplemental Figure S10).

S-acylation was shown to promote the segregation of the transmembrane protein Linker for Activation of T cells (LAT) into membrane domains in giant PM vesicles isolated from animal cells (Levental et al., 2010). In addition, S-acylation drives the association of the membrane protein spike from SARS-CoV-2 with ordered lipid domains (Mesquita et al., 2021). However, in plants, the role of S-acylation in the targeting of proteins into nanodomains remains elusive (Jaillais and Ott, 2020). Interestingly, the S-acylation of remorins was shown to be required for their PM targeting but not for their organization in nanodomains (Konrad et al., 2014). Here, we demonstrated that the N-terminal part of HIR2 carrying the S-acylation sites can drive a GFP to the PM but does not induce its clustering in nanodomains (Figure 7), showing that the S-acylation of HIR2 is not sufficient for its nanodomain organization. However, since HIR2^C6A^ and HIR2^C7A^ are normally S-acylated and because HIR2^C6A, C7A^ is retained intracellularly, it is difficult to conclude on a putative contribution of S-acylation to HIR2 clustering process in the PM.

### Plasma membrane lipid composition influences HIR2 organization

We investigated the role of lipids in the organization of HIR2 using pharmacological treatments altering PM lipid composition combined with microscopy analyses. We showed using TIRF microscopy, that proper sterol composition of the PM is required for the nanoclustering of HIR2 in the PM of Arabidopsis root epidermal cells (Figure 5). This result is in accordance with a role of sterols in the PM dynamics of another HIR isoform, HIR1, that was previously reported, using the sterol-depleting agent mβCD (Lv et al., 2017). Our results reinforce the notion that in plant cells, sterols play a key role in protein nano-organization as highlighted in the past (Gronnier et al., 2017; Grison et al., 2019; Cao et al., 2020). The reduction of HIR2 clustering following fenpropimorph treatment was not accompanied by a higher mobility of HIR2 molecules in the PM, as revealed by sptPALM (Figure 5). Therefore, nanodomain organization and low lateral mobility of proteins are not necessarily correlated. VLCFA are essential lipids that play a structural role in the PM (Batsale et al., 2021). Altering the composition of VLCFA dramatically reduced the clustering of HIR2 in the PM of Arabidopsis root epidermal cells which was accompanied by a strong increase in the mobility of HIR2 molecules (Figure 5), arguing for an important role of VLCFA in HIR2 nanodomain organization. VLCFA-containing lipids include sphingolipids and phosphatidylserine. So far, our pharmacological approach does not allow us to identify precisely which lipid(s) bearing a VLCFA is/are involved in HIR2 nano-organization. Phosphatidylserine constitutes a very interesting candidate since this lipid can arrange in nanoclusters and was shown to stabilize ROP6 protein in PM nanodomains (Platre et al., 2019).

### The oligomerization of HIR2 through its C-terminal part determines the organization of HIR2 into PM nanodomains, which is required for its proper function

We sought to identify the domains involved in HIR2 homo-oligomerization and highlighted that the C-terminal part of HIR2 is required for self-assembly. Despite the very low level of amino acid identity commonly found between the members of the SPFH superfamily, we showed that a partially conserved hydrophobic motif present in both the very C-terminal parts of HIR2 and human Stomatin plays a crucial role in oligomerization (Figure 6) (Umlauf et al., 2006). However, since the deletion of the whole C-terminal part of HIR2 had even stronger deleterious effect on HIR2 oligomerization, we can assume that several C-terminal zones are implicated in HIR2 self-assembly. Our results in combination with other works reinforce the prominent function of the C-terminal part of SPFH proteins in their oligomerization (Umlauf et al., 2006; Solis et al., 2007; Yu et al., 2017). However, HIR2 protein may carry additional oligomerization domains and recently, valine 157 present in the SPFH domain of Arabidopsis HIR4 isoform, which is also conserved in HIR2, was shown to be important for HIR4 oligomerization (Liu et al., 2024). Although the exact oligomeric state of HIR2 remains to be determined, previous studies suggested that HIR2 protein may exist as dimers, trimers and tetramers (Qi et al., 2011). However, since other SPFH proteins form very large oligomers, 12 copies of HflK–HflC dimers in bacteria and 22 pairs of Flot1 and Flot2 subunits in humans (Ma et al., 2022; Fu and MacKinnon, 2024), we suspect that HIR2 oligomeric state is higher than a tetramer. We showed that the C-terminal part of HIR2 is dispensable for PM targeting and that indeed the 17 first amino acids of HIR2 were sufficient to drive a fluorescent protein to the PM (Figure 7). However, the C-terminal part of HIR2 is crucial for the organization of HIR2 in PM nanodomains since: (i) sequential deletions of the C-terminal part attenuated (HIR2^1-254^) or totally abolished HIR2 nanoclustering (HIR2^1-182^), (ii) the C-terminal part of HIR2 artificially targeted to the PM arranged into clusters (HIR2^Nter-Cter^, Figure 7). Given the role of the C-terminal part of HIR2 in its self-assembly, we propose that HIR2 oligomerization is necessary and sufficient for the organization of HIR2 into PM nanodomains in plant cells. A crucial role for oligomerization in the arrangement into nanodomains could be probably extended to other plant proteins such as remorins. Indeed, the formation of high-order complexes constituted of remorin trimers was previously proposed to play a role in remorin organization into membrane nanodomains (Bariola et al., 2004; Perraki et al., 2012; Legrand et al., 2019; Martinez et al., 2019), although to our knowledge, formal evidences from plant cell biology experiments are still missing. sptPALM analysis revealed that abolishing HIR2 nanoclustering trough a deletion of the C-terminal part considerably increase the lateral mobility of HIR2 single molecules in the PM (Figure 7). This result is in accordance with data showing that mutated human Stomatin proteins that cannot oligomerize any more display an increase in their lateral mobility in the PM, as revealed by fluorescence recovery after photobleaching, suggesting a conserved mechanism between plant and animals (Umlauf et al., 2006).

HIR2 protein is a positive actor of plant immunity, although the underlying molecular mechanisms remain largely unknown. Here, we showed that HIR2 overexpression stimulated apoplastic ROS accumulation in plant cells in response to flg22, a mechanism that required HIR2 oligomerization and nanodomain organization in the PM (Figure 8). This result is in line with a recent work showing that the oligomerization of Arabidopsis HIR4 was essential for its role in mediating immunity via a regulation of ROS burst (Liu et al., 2024). Interestingly, pathogens can interfere with HIR oligomerization via effectors to dampen the plant immune response (Liu et al., 2024). Taking into consideration our results, we can assume that this mechanism occurs through a loss of HIR nanodomain organization in the PM. SPFH proteins were proposed to act as scaffolding proteins in membrane nanodomains (Langhorst et al., 2005), although experimental evidences remained limited so far. Our data showing that HIR2 possesses the intrinsic property, through oligomerization, to arrange in nanoclusters support this concept of nanodomain structuration ensured by SPFH proteins. HIR2 may interact with and recruit proteins in specific nanodomains, such as some proteins involved in plant immunity that were shown to physically interact with HIR (Jung and Hwang, 2007; Zhou et al., 2009; Qi et al., 2011; Lv et al., 2017). In the future, it will be interesting to determine how HIR-containing nanodomains may act as hubs implicated in the defense against pathogens, notably by influencing ROS production. We showed that HIR2 defines nanodomains mostly distinct from those labeled by the remorins REM1.2 and REM1.3 in Arabidopsis cells (Figure 1). Although HIR2 and remorins are both implicated in plant immunity (Raffaele et al., 2009; Qi et al., 2011), their spatial segregation in distinct PM nanodomains suggest that they act in different pathways. Intriguingly, a partial co-localization in PM nanodomains between HIR1 and REM1.3 was reported in Arabidopsis leaf epidermal cells (Lv et al., 2017). Since the nanodomain patterning in the PM is extremely complex and proteins from the same family can organize in different nanodomains (Jarsch et al., 2014), it is tempting to speculate that HIR1 and HIR2 proteins may arrange in distinct nanodomains in the PM of Arabidopsis cells.

A long standing and complex question in the field of membrane nanodomains is whether proteins are targeted to pre-existing lipid-mediated nanodomains or if they actively organize their lipid environment through protein-lipid interactions (Gouguet et al., 2021). Interestingly, overexpression of remorins in plant cells modify the local membrane order (Huang et al., 2019), suggesting that these proteins, which oligomerize and interact with lipids (Perraki et al., 2012; Gronnier et al., 2017), can influence their lipid environment. In addition, Legrand and coworkers showed by atomic force microscopy in model membranes that *Solanum tuberosum* StREM1.3 can organize lipid nanodomains, even in absence of pre-existing lipid domains (Legrand et al., 2023). Although not directly related to nanodomain organization, remorins were recently shown to stabilize membrane topologies in a cell wall independent manner (Su et al., 2023). By forming large multimeric complexes and binding to specific lipids, animal SPFH proteins were hypothesized to actively form nanodomains enriched in certain lipids (Browman et al., 2007). Since HIR2 actively participates in the formation of nanodomains and probably interacts with lipids as stated above, we anticipate that HIR2 may play a role in lipid clustering, although experimental evidences are needed. Recently, original works on SPFH proteins from bacteria and human highlighted that these proteins assemble in very large circular complexes that delimit a membrane domain of about 20-30 nm in diameter, which could de facto allows the segregation of lipids but also proteins (Ma et al., 2022; Fu and MacKinnon, 2024).

## MATERIAL AND METHODS

### Genetic constructions

All the constructs described in this section were obtained using the MultiSite Gateway® Three-Fragment Vector Construction system (Life Technologies). The primer sequences are indicated in Supplemental Table 1. The *HIR2* promoter corresponding to a sequence of 1,241 pb upstream of the *HIR2* start codon was amplified from *Arabidopsis thaliana* genomic DNA using the Fw1 forward and Rv1 reverse primers and subsequently cloned into the pDONR.P4P1R entry vector. The *HIR2* open reading frame (ORF) without the stop codon was amplified from *Arabidopsis* cDNAs with the Fw2 and Rv2 primers and cloned into the pDONR.221 vector. Site directed mutagenesis of HIR2 ORF was performed on the pDONR.221-HIR2 vector by the SPRINP method (Edelheit et al., 2009), using the primers Fw3 – Fw7 and Rv3 – Rv7 to generate *HIR2^G2A^*, *HIR2^C6A^*, *HIR2^C7A,^ HIR2^C58A^, HIR2^C6, C7A^, HIR2 ^G2A, C6A, C7A,^ HIR2 ^C6A, C58A^, HIR2 ^C7A, C58A^ and HIR2 ^C6A, C7A, C58A^*. Repeated mutation steps were applied when necessary. Truncated versions of HIR2 coding sequence were amplified with primers Fw2 and Rv8 or Rv9 to obtain *HIR2^1-182^* (codons 1-182) or *HIR2^1-254^* (codons 1-254), respectively. The 17 first codons of *HIR2* (*HIR2^Nter^*) were joined with mEGFP coding sequence by PCR with primers Fw8 and Rv10 and introduced into pDONR.221. *HIR2^Cter^*was generated by amplifying the last 103 codons of *HIR2* without the stop codon with Fw9 primer, that allowed adding a start codon, and Rv2 primer. The chimeric sequence *HIR2^Nter-Cter^* was obtained by fusing by PCR the 17 first codons of *HIR2* to the last 103 codons of *HIR2* without the stop codon with Fw10 and Rv2 primers. *HIR2^1-182^*, *HIR2^1-254^*, *HIR2^Nter^-mEGFP*, *HIR2^Cter^* and *HIR2^Nter-Cter^* sequences were subsequently cloned into pDONR.221. The coding sequences of mEOS2 and mEGFP with stop codons were amplified with Fw11/Rv11 and Fw12/Rv12 primers, respectively and cloned into the pDONR.P2RP3 entry vector to allow C-terminal fusions. Entry vectors carrying the *p35S* promoter (pDONR.P4P1R-p35S), the *AHA2* ORF (pDONR.221-AHA2) or mCherry coding sequence allowing C-terminal fusions (pDONR.P2RP3-mCherry) were previously described (Marques-Bueno et al., 2016; Martin-Barranco et al., 2020). Final destination vectors for expression in plants were obtained by multisite Gateway® recombination using the entry vectors described above and the pH7m34GW and pK7m34GW destinations vectors used for mCherry and EGFP/mEOS2 fusions, respectively. The following constructs were generated: pHIR2::HIR2-mCherry/mEGFP, pHIR2::HIR2^G2A^-mCherry, pHIR2::HIR2^C6A^-mCherry, pHIR2::HIR2^C7A^-mCherry, pHIR2::HIR2^C58A^-mCherry, pHIR2::HIR2^C6A, C7A^-mCherry, pHIR2::HIR2^G2A, C6A, C7A^-mCherry, pHIR2::HIR2^C6A, C58A^-mCherry, pHIR2::HIR2^C7A, C58A^-mCherry, pHIR2::HIR2^C6A, C7A, C58A^-mCherry, pHIR2::HIR2^1-182^-mCherry/mEGFP, pHIR2::HIR2^1-254^-mCherry/mEGFP, pHIR2::HIR2^Nter^-mEGFP, pHIR2::HIR2^Cter^–mCherry, pHIR2::HIR2^Nter-Cter^-mCherry, pHIR2::HIR2-mEOS2, pHIR2::HIR2^1-182^-mEOS2, p35S::HIR2-mEGFP, p35S::AHA2-mCherry. p35S::Sar1(DN)-YFP and p35S::HDEL-pHluorine vectors were previously described (daSilva et al., 2004; Moreau et al., 2021).

### Arabidopsis transgenic lines and growth conditions

*Arabidopsis thaliana* Col-0 ecotype was transformed with the different genetic constructions described above by the floral dipping technique using *Agrobacterium tumefaciens* GV3101 strain. Arabidopsis transgenic line expressing *35S::GFP-StREM1.3* was previously described (Legrand et al., 2023). Arabidopsis transgenic lines co-expressing *35S::GFP-StREM1.3*, *pREM1.2::YFP-REM1.2* or *pREM1.3::YFP-REM1.3* (Jarsch et al., 2014) with *pHIR2::HIR2-mCherry* were obtained by crossing. Arabidopsis plants were vertically grown in sterile conditions at 21°C with 16 h light/8 h dark cycles with a light intensity of 90 µmol m^-2^ s^-1^ using Philips 17W F17T8/TL741 bulbs. The plant growth medium used was half-strength Murashige and Skoog (MS/2) medium containing 1% sucrose and 1% agar.

### Transient expression of proteins in *N. benthamiana* leaves

Leaves from 4-week old plants of *Nicotiana benthamiana* were agroinfiltrated with *Agrobacterium tumefaciens* strain GV3101 suspensions of optical density 0.6 - 0.9, as previously described (Voinnet et al., 2003). After 2 to 3 days, infected leaves were collected for microscopy and/or biochemistry analysis.

### Pharmacological treatments

Nine-day old Arabidopsis seedlings grown on MS/2 solid medium were transplanted onto plates with the same solid medium supplemented with 2-bromopalmitate (10 µM) for 7 hours, fenpropimorph (50 µg / mL) for 24 hours (Gronnier et al., 2017), or metazachlor (100 nM) for 48 hours (Grison et al., 2019). Since, these drugs were diluted 4,000 times from stock solutions in DMSO, 4,000 times diluted DMSO was used as mock treatment.

### Co-immunopurifications

Co-Immunopurifications (co-IP) were performed on approximately 500 mg of *N. benthamiana* leaves transiently co-expressing HIR2-mCherry with HIR2-mEGFP, HIR2^1-182^-mEGFP or HIR2^1-254^-mEGFP proteins, as previously described (Martin-Barranco et al., 2020). Briefly, tissues were ground in liquid nitrogen and resuspended in solubilization buffer (50 mM Tris-HCl (pH 7.4), 150 mM NaCl, 5 mM EDTA, 1% n-Dodecyl β-D-maltoside (DDM) and plant specific protease inhibitors (Sigma-Aldrich)). After two successive centrifugation steps at 3,800 × *g* for

10 min at 4 °C, the resultant supernatants were collected and solubilization of membrane proteins was continued for 1 h at 4 °C on a rotating wheel. Samples were then centrifuged at 100,000 × *g* for 1 h at 4 °C to remove unsolubilized material and supernatants containing solubilized proteins were recovered for co-IPs. HIR2, HIR2^1-182^ and HIR2^1-254^ proteins fused to mEGFP were immunopurified using µMACS GFP isolation kit (Miltenyi Biotec), following the instructions of the manufacturers. Before elution, extensive washes were performed with solubilization buffer. Immunodetections of GFP and mCherry fusion proteins were performed as described below. Co-IP combined with immunodetections were performed three times with similar results.

### Immunopurification of HIR2-mCherry for mass spectrometry analysis

Immunopurification of HIR2-mCherry was performed on approximately 1 g of 10-day-old Arabidopsis plantlets expressing HIR2-mCherry under the control of *HIR2* promoter. Protein solubilization was performed as described above using 1% DDM. Following an ultracentrifugation at 100,000 × *g* for 1 h at 4 °C, solubilized proteins were recovered and mCherry fusion protein was immunopurified using RFP-Trap®_MA magnetic beads (Chromotek), following the instructions of the manufacturers, as previously described (Martin-Barranco et al., 2020). Before elution, extensive washes were performed with solubilization buffer. Immunodetection of HIR2-mCherry was performed as described below.

### Mass spectrometry: sample preparation, analysis and identification

Immunopurified HIR2-mCherry was loaded on a 10 % precast gel (Biorad) and submitted to a 15 min migration at 100 V for a 1 cm migration. The lane was cut into small pieces and processed as described (Berger et al., 2022). Briefly, gel pieces were washed twice with carbonate buffer (25 mM NH4HCO3), then with 50 % acetonitrile in 25 mM NH4HCO3 before being fully dehydrated with 100% acetonitrile and dried at room temperature. After protein reduction for 45 min at 56 °C with 10 mM DTT and then alkylation with 55 mM iodoacetamide for 30 min at room temperature in the dark, gel pieces were washed twice with 50 % acetonitrile in 25 mM NH4HCO3 and then dehydrated in 100 % acetonitrile. The sample was digested with 188 µg trypsin (Sequencing Grade Modified Trypsin, Promega) in 25 mM NH4HCO3 overnight at 37 °C. Peptides were first extracted from the gel with 50 µl of 2 % formic acid (FA) and then twice with 80 % acetonitrile / 2 % FA. Supernatants were pooled and vacuum-dried, to be resuspended in 8 μl of 2 % FA.

Peptides were analyzed via liquid-chromatography tandem mass spectrometry (LC-MS/MS) using standard settings as described (Berger et al., 2022) using a Ultimate 3000 RSLC nano system (Thermo Fisher Scientific) as an HPLC system and a Exploris 240 Plus Orbitrap mass spectrometer (Thermo Fisher Scientific) as a mass analyser. Peptides were identified via MS/MS based on the information-dependent acquisition of fragmentation spectra of multiple charged peptides. A data-dependent acquisition MS method was used with resolutions of 120 000 for MS and 15 000 for MS2. Spectrum were recorded with Xcalibur software (4.4.16.14) (Thermo Fisher Scientific).

Mass spectrometry raw data were processed using the Proteome Discoverer software (Version 1.4.0.288, Thermo Fisher Scientific, Bremen, Germany) and Mascot (version.2.4, Matrix Science, http://www.matrixscience.com) as search engine. The MS data were matched against Tair10 implemented with HIR2-mCherry sequence. Carbamidomethylation of cysteine was set as a fixed modification, and the oxidized methionine (M), acetylation (protein N-term) deamidation of asparagine and glutamine, N-terminal-pyroglutamylation of glutamine and glutamate, oxidation of methionine and myristoylation (N-terminal G) were set as variable modifications. Trypsin was specified as the digesting protease, and up to two missed cleavages were allowed. The mass tolerance for the database search was set to 10 ppm for full scans and 0.02 Da for fragment ions.

### Cell fractionation

Cell fractionation was performed on 200 mg of *N. benthamiana* leaves transiently expressing WT HIR2 and HIR2 mutated proteins fused to mCherry. Tissues were ground in liquid nitrogen and resuspended in 400 µl of microsomal extraction buffer (500 mm sucrose, 50 mm HEPES (pH 7.4), 1 mm DTT, 5 mm EDTA, 5 mm EGTA and plant specific protease inhibitors (Sigma-Aldrich)). Extracts were centrifuged at 13,000 × *g* for 20 min at 4 °C to eliminate cell debris. Supernatants were centrifuged again at 13,000 × *g* for 10 min at 4 °C. Supernatants were then subjected to ultracentrifugation at 100,000 × *g* for 90 min at 4 °C. Supernatants were collected (cytosolic fractions) and pellets were resuspended in an equal volume of microsomal extraction buffer (microsomal fractions).

### PEG shift assay

The S-acylation of HIR2 and HIR2 mutated proteins was analyzed through a PEG shift assay adapted from the protocol developed by Percher and co-workers (Percher et al., 2017). Briefly, 150 mg of ten-day-old Arabidopsis plantlets expressing wild-type HIR2 and point mutated HIR2 fused to mCherry under the control of *HIR2* promoter were ground in liquid nitrogen. Tissues were resuspended in lysis buffer (TEA buffer (50 mM triethanolamine, 150 mM NaCl, pH7.3) containing 4% SDS, 5 mM EDTA and plant specific protease inhibitors (Sigma-Aldrich)). After centrifugation at 3,800 × *g* for 10 min at 4°C, the supernatants were recovered and aliquoted by 185 µL. To ensure reduction of protein disulfide bonds, 10 μL of 200 mM Tris-(2-carboxyethyl)-phosphine pH 7.3 were added to protein samples that were then mixed for 30 min at room temperature. Then, reduced cysteines were capped by adding 5 μL of 1 M N-ethylmaleimide (NEM). Samples were mixed for 2 h at room temperature. Subsequently, proteins were precipitated by sequentially adding 800 µl of methanol, 300 µl of chloroform and 600 µl of H2O, at 4°C. After mixing, samples were centrifuged at 20,000 × *g* for 5 min at 4°C, the aqueous (top) phase was removed and 1 ml of pre-chilled methanol was added. After mixing and a centrifugation at 20,000 × *g* for 3 min at 4°C, the protein pellet was washed with 800 μL of pre-chilled methanol. After centrifugation at 20,000 × *g* for 3 min at 4°C, the supernatant was discarded and the protein pellet was dried. To ensure complete removal of NEM, this precipitation step was repeated once after resuspending the protein pellet in 200 µl of TEA buffer pH 7.3, 4% SDS. Then, dry protein pellets were resuspended in 60 μL of TEA buffer containing 4% SDS and 4 mM EDTA, and each sample was divided into two 30 μL for the +/− hydroxylamine treatments. S-acyl groups coupled to cysteine via a thioester bond were cleaved by adding 90 μL of 1 M hydroxylamine, pH 7.4. A control without hydroxylamine was performed by adding 90 μL of 50 mM Tris HCl, pH 7.4. Samples were mixed for 1 h at room temperature. Then, to remove hydroxylamine, proteins were precipitated using a mix of methanol, chloroform and H2O, as described above. The dry pellet of proteins was resuspended in 30 μL of TEA buffer containing 4% SDS and 4 mM EDTA before adding 90 μL of TEA buffer pH 7.3 containing 0.2 % Triton X100 and 1.33 mM methoxy polyethylene glycol maleimide (mPEG-Mal) with a molecular weight of 5 kDa (Sigma, 63187). Samples were mixed for 2 h at room temperature. During this step, exposed cysteines are labeled with mPEG-mal at stoichiometries and levels reflective of the original number of S-fatty acylation events (Percher et al., 2017). Then, proteins were precipitated using a mix of methanol, chloroform and H2O, as described above. Dry protein pellets were resuspended in 30 μL of 2x Laemmli buffer (2.5 % SDS, 20% glycerol (v/v), 4% dithiothreitol, 0.01 % bromophenol blue, 125 mM Tris HCl pH 6.8) and heated for 5 min at 95 °C. Immunodetection of mCherry fusion proteins was performed as described below.

### Immunodetections

Immunoblot analyses were performed as previously described (Barberon et al., 2011). Immunodetections of GFP and mCherry fusion proteins were performed using an anti-GFP antibody conjugated to horseradish peroxidase (HRP) (Miltenyi Biotec 130-091-833, 1/5,000) and a rabbit anti-DsRed antibody (Clontech 632496, 1/5,000), respectively. For cell fractionation experiments, H^+^-ATPases and cytosolic fructose-1,6-bisphosphatase (FBPase) were immunodetected using a rabbit antibody raised against the Plasma Membrane H^+^-ATPase 2 (PMA2) from *Nicotiana plumbaginifolia* (W1C) diluted 1/15,000 (Morsomme et al., 1998) and a rabbit anti-FBPase antibody (Agrisera AS04043), diluted 1/10,000, respectively. The anti-rabbit IgG secondary antibody coupled to HRP was diluted 1/20,000. Detection of HRP chemiluminescence was performed using SuperSignal West Dura Extended Duration Substrate (Thermo Scientific) in a Chemidoc Touch Imaging system (Bio-Rad).

### Measurement of apoplastic ROS in *N. benthamiana* leaf discs

Measurements of apoplastic ROS production following flg22 treatment in *N. benthamiana* leaf disks transiently expressing HIR2-mCherry, HIR2^1-182^-mCherry and mCherry under the control of *HIR2* promoter were performed as follows, mainly as described by D’Ambrosio and coworkers (D’Ambrosio et al., 2017). Two days post infiltration, expression of mCherry fusion proteins and mCherry alone was verified by confocal microscopy as described below. Then, leaf discs of 4 mm in diameter were generated with a cookie cutter and incubated for 24 hours in 96-well plates containing sterile S Buffer (10 mM MES-KOH pH 6.1 mM CaCl2, 1 mM KCl and 1% sucrose). Afterwards, the discs were washed 3 times with R buffer (10 mM HEPES-KOH pH 8) and then incubated with T buffer (10mM HEPES-KOH pH 8, 20 mM Luminol L-012 (Sigma) and 10 mM Horseradish Peroxidase (Sigma). Subsequently, apoplastic ROS production of leaf discs treated or not with 100 nM flg22 was evaluated by measuring the luminescence generated by the oxidation of the luminol in presence of ROS, for 80 min, using a CLARIOstar plate reader machine (BMG-Lab tech). For normalization, mCherry fluorescence of all discs was measured at 614 nm with the CLARIOstar after excitation at 580 nm.

### Confocal and spinning-disk microscopy, image analysis

Entire seedlings were mounted in a drop of liquid medium of the same composition as the medium used for a given treatment. A laser scanning microscope (SP8, Leica) equipped with a HyD detector or a spinning disk microscope (Dragonfly, Andor) equipped with EMCCD iXon888 camera (Andor) were used for imaging. 40x Plan Apo (N. A. 1.1, water immersion), and 100x Plan Apo (N.A. 1.45, oil immersion) objectives were used with the laser scanning microscope, and the spinning disk microscope, respectively. 488 nm, and 561 nm laser lines were used for mEGFP or YFP, and mCherry excitation, respectively.

Following spinning-disk acquisitions, nanodomain co-localization was evaluated with JACoP plugin (Bolte and Cordelieres, 2006) in Fiji (Schindelin et al., 2012) on square regions of interest (ROIs) of 5 times 5 µm. Rotation of one channel by 90° with respect to the other was performed as a control for random co-localization.

Coefficient of variance (CV) of signal intensities (Retzer et al., 2017) in linear probe of 10 µm was determined in order to assess the level of clustering as in (Danek et al., 2020).

Maximum intensity projections, average intensity projections and orthogonal projections were constructed in Fiji.

### TIRF microscopy

TIRF microscopy was performed using the same inverted Zeiss microscope as for sptPALM (see the next section). One hundred images were acquired with 50 ms exposure time and the electronic gain set to 300, with 488 nm excitation (OBIS LX 50 mW; Coherent) and a 525/22.5 nm emission filter (Chroma) for mEGFP, and 561 nm excitation (SAPPHIRE 100 mW; Coherent) and a 600/50 nm emission filter (Chroma) for mCherry. An average intensity projection was computed from the 100 acquired images. CV values were calculated as described above.

### Single-particle tracking photoactivated localization microscopy

The sptPALM experiments were conducted following the protocol outlined by (Bayle et al., 2021). Briefly, observations of Arabidopsis root epidermal cells and *N. benthamiana* leaf epidermal cells were conducted utilizing a custom-built TIRF microscope featuring an electron-multiplying charge-coupled device camera (Andor iXON XU_897) and a 100× oil plan-apochromat objective (Zeiss) with a numerical aperture of 1.46. Pixel size was 102 nm. Manual adjustment of the laser angle ensured optimal generation of evanescent waves, maximizing the signal-to-noise ratio during imaging of the cell membrane in close proximity to the coverslip. Photoconversion of the mEOS2 tag was achieved through low-intensity illumination at 405 nm (OXXIUS LBX-405 set at 40mW; Coherent), while image acquisition utilized 561 nm excitation (SAPPHIRE 100 mW; Coherent) combined with a 600/50 emission filter (Chroma). A total of four thousand images were captured per region of interest (ROI) and streamed into open source acquisition software (Barho et al., 2022) at a 30 ms exposure time. Analysis was conducted on ten to twenty cells per replicate, with three biological replicates.

### Single-particle tracking and Vonoroï tessellation

Single molecules were individually localized and tracked using the ImageJ pluggin Trackmate 7 software (Ershov et al., 2022). Subsequently, dynamic characteristics of these single emitters within cells were deduced from their tracks using a custom-made analysis software developed in MATLAB (The MathWorks) (Bayle et al., 2021). The diffusion coefficient (D) for each track was computed by fitting the mean squared displacement (MSD) curve accordingly (Bayle et al., 2021). For clustering analysis, the positions provided by Trackmate 7 for each mEOS2 detection served as inputs for the SR-Tesseler software which in turn was used to generate Vonoroï diagram (Levet et al., 2015). Corrections for multiple detections of the same emitter were implemented based on recommendations from the same reference. The local densities of each track were computed as the inverse of their minimal surface.

### Statistical analysis

For microscopy experiments, a representative image is shown. Statistical analyses were performed using the software GraphPad Prism 7. The sample size and statistical tests used are mentioned in the figure legends.

## ACKNOWLEDGMENTS

We thank Yvon Jaillais and Lionel Verdoucq for interesting scientific discussions. We also thank Marc Boutry and Nadine Paris for the anti-PMA2 antibody and p35S::HDEL-pHluorine vector, respectively. This work benefited from access to the optic facilities of the Integrated Biophysics and Structural Biology Platform (PIBBS). PIBBS is a GIS-IBISA platform and belongs to the French Infrastructure for France Bio-Imaging (FBI), supported by the National Research Agency (ANR-10-INBS-04-01). Mass spectrometry experiments were carried out using the facilities of the Montpellier Proteomics Platform (PPM, BioCampus Montpellier). We thank the Montpellier Ressources Imagerie (MRI) and the Histocytology and Plant Cell Imaging Platform (PHIV) for providing the microscope facility.

## SUPPLEMENTAL DATA

**Supplemental Figure S1.**
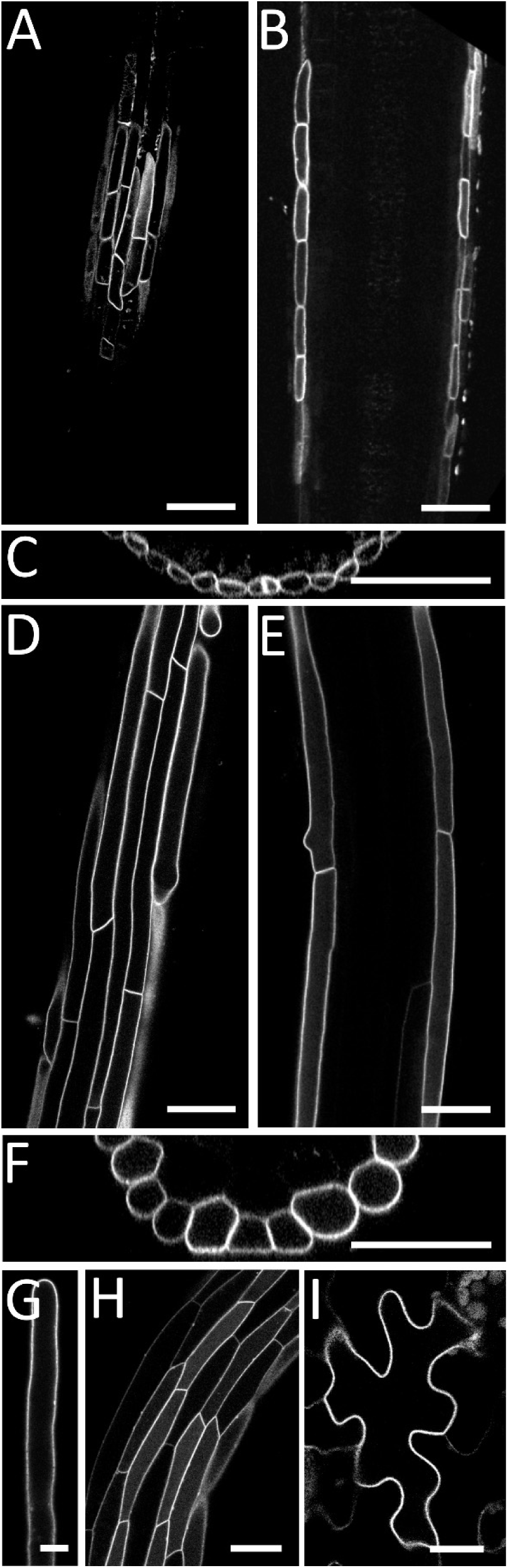
Expression territories and subcellular localization of HIR2 in *Arabidopsis thaliana*. 11-day-old plants expressing HIR2-mCherry fusion protein under the control of *HIR2* promoter were imaged by scanning confocal microscopy. HIR2-mCherry signal was observed in the PM in lateral root cap cells **(A-C)**, in epidermal cells from the root differentiated zone **(D-F)**, in root hairs (**G**), in hypocotyl cells **(H)** and in cotyledon epidermal pavement cells **(I)**. (**A, D, G, H, I):** Secant view of the cells. **(B)** and **(E)**: Tangential view in the middle of the root. **(C)** and **(F)**: XZ orthogonal projection from the Z-stack corresponding to A, B, and D, E, respectively. Scale bars represent 50 µm (A, B, D, E, H), 20 µm (C, F, I) or 10 µm (G).

**Supplemental Figure S2.**
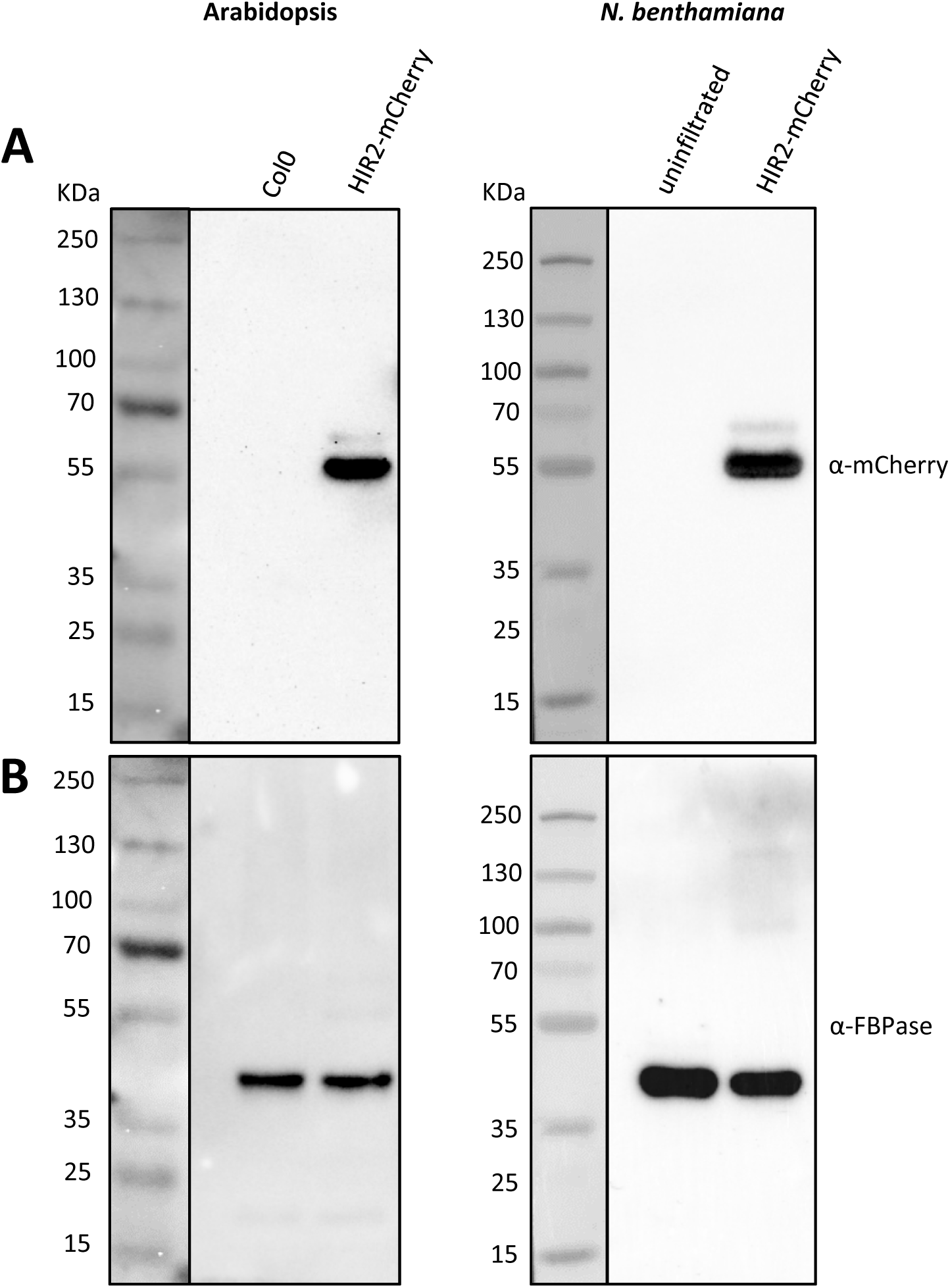
The anti-mCherry antibody specifically recognizes HIR2-mCherry fusion protein in plant protein extracts. **(A)** Anti-mCherry immunoblots were performed on total protein extracts from: WT Arabidopsis (Col0) and Arabidopsis seedlings expressing HIR2-mCherry (left), uninfiltrated *N. benthamiana* and *N. benthamiana* transiently expressing HIR2-mCherry (right). In both Arabidopsis and *N. benthamiana,* HIR2-mCherry was expressed under the control of *HIR2* promoter. Arabidopsis plantlets were grown 11 days on MS/2 medium and tobacco leaves were collected 2 days after infiltration. The expected molecular weight of HIR2-mCherry is 58.9 kDa. **(B)** After stripping, the transfer membranes were re-probed with an antibody recognizing the cytosolic FBPase used as a protein loading control. In (A) and (B), the protein ladder transferred on the membranes is shown on the left of the immunoblots.

**Supplemental Figure S3.**
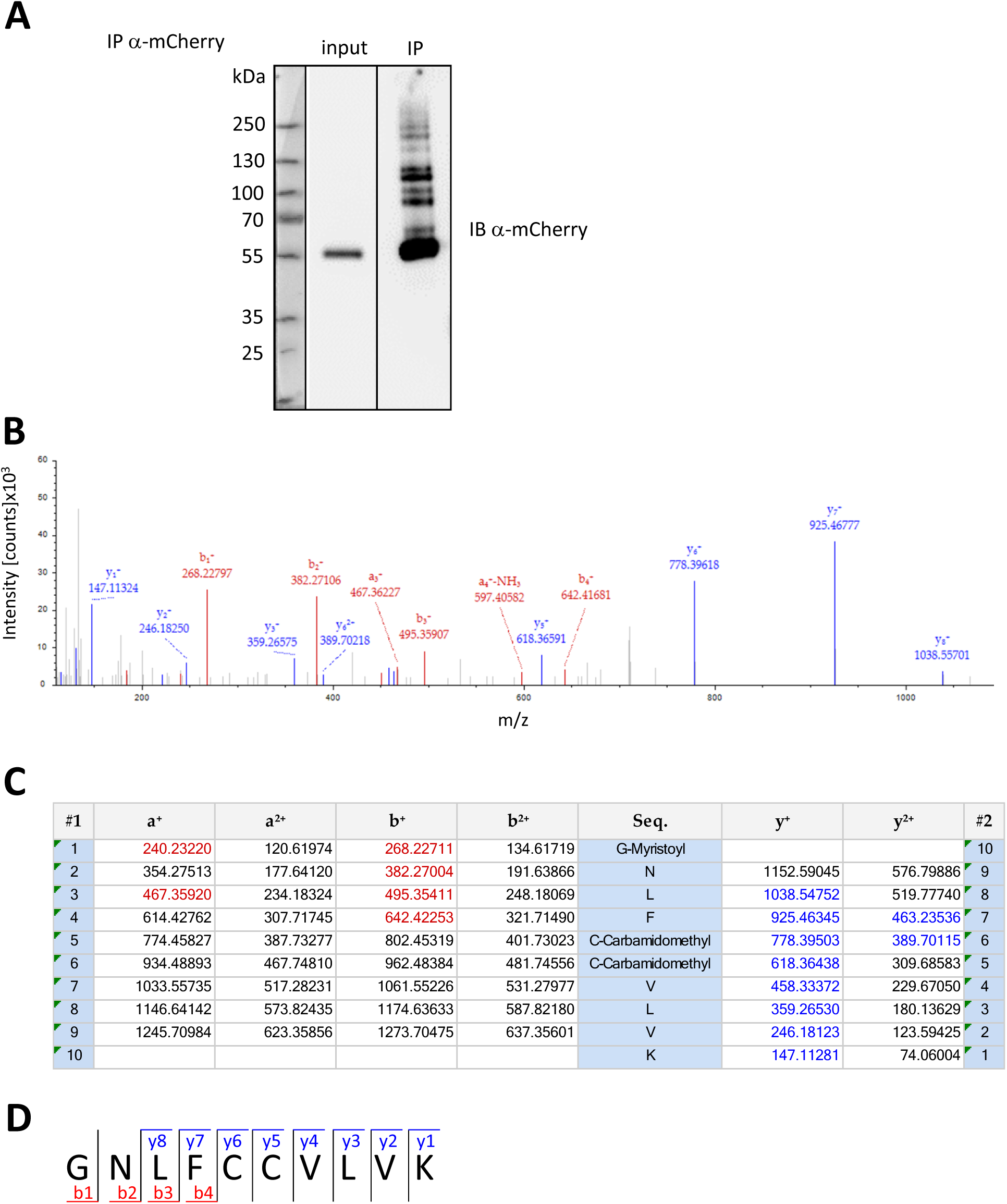
Identification of HIR2 N-terminal myristoylation. **(A)** HIR2-mCherry expressed under the control of *HIR2* promoter in Arabidopsis plantlets was immunopurified (IP) from root solubilized proteins using an anti-mCherry antibody coupled to magnetic beads, then, immunoblotting (IB) with an anti-mCherry antibody was performed on inputs and IP fractions. **(B)** Tandem mass spectrometry (MS/MS) spectrum corresponding to the fragmentation of a N-terminal peptide of HIR2-mCherry immunopurified fusion protein, with ion score Mascot 44 and sequence GNLFCCVLVK. (**C)** Table of identified masses. Fixed modification (carbamidomethylation, 57.02146 Da ) and variable modification (myristoylation, 210.19837 Da) were considered. N-terminal and C-terminal fragments are indicated in red and blue characters, respectively (B, C). **(D)** Schematic representation of peptide fragmentation. Peptide characteristics are: Charge: +2, monoisotopic m/z: 710.41046 Da, MH^+^: 1419.81365 Da, retention time: 132.74 min.

**Supplemental Figure S4.**
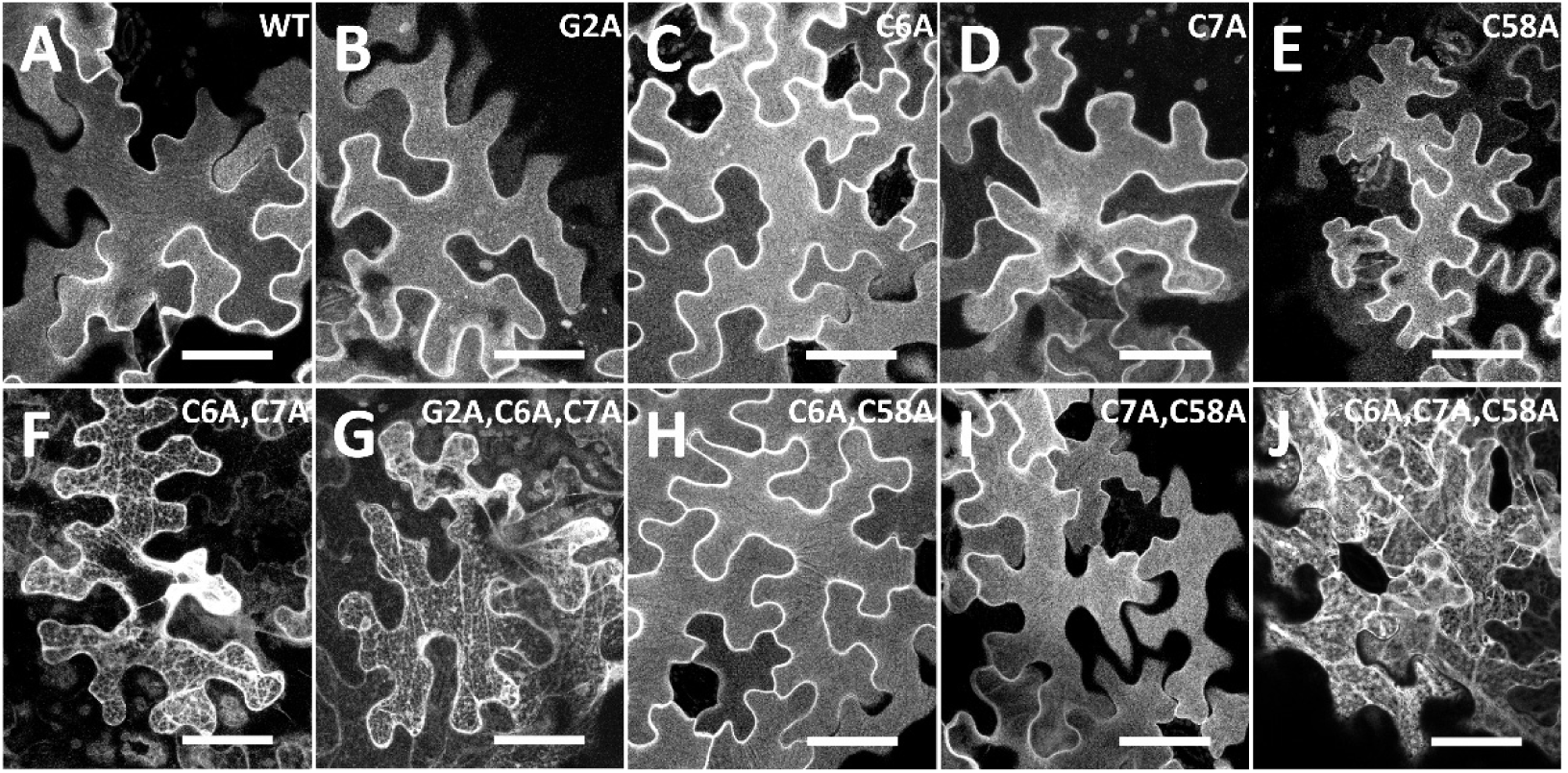
Simultaneous mutations of C6 and C7 induce HIR2 intracellular retention. Maximum intensity projections constructed from Z-stack of 25 focal planes acquired by scanning confocal microscopy on *N. benthamiana* leaf epidermal cells expressing HIR2 point mutants fused to mCherry under the control of *HIR2* promoter. **(A)**: HIR2-mCherry, **(B)** HIR2^G2A^-mCherry, **(C)**: HIR2^C6A^-mCherry, **(D)**: HIR2^C7A^-mCherry, **(E)**: HIR2^C58A^-mCherry, **(F)**: HIR2^C6A, C7A^-mCherry, **(G)**: HIR2^G2A, C6A, C7A^-mCherry, **(H)**: HIR2^C6A, C58A^-mCherry, **(I)**: HIR2^C7A, C58A^-mCherry, **(J)**: HIR2^C6A, C7A, C58A^-mCherry. Note that the PM localization of HIR2 and its variants in A-E, H, I contrasts with the intracellular localization observed in F, G, J. Maximum intensity projections were constructed from the same cells as in Figure 2A-J. Scale bars represent 50 µm.

**Supplemental Figure S5.**
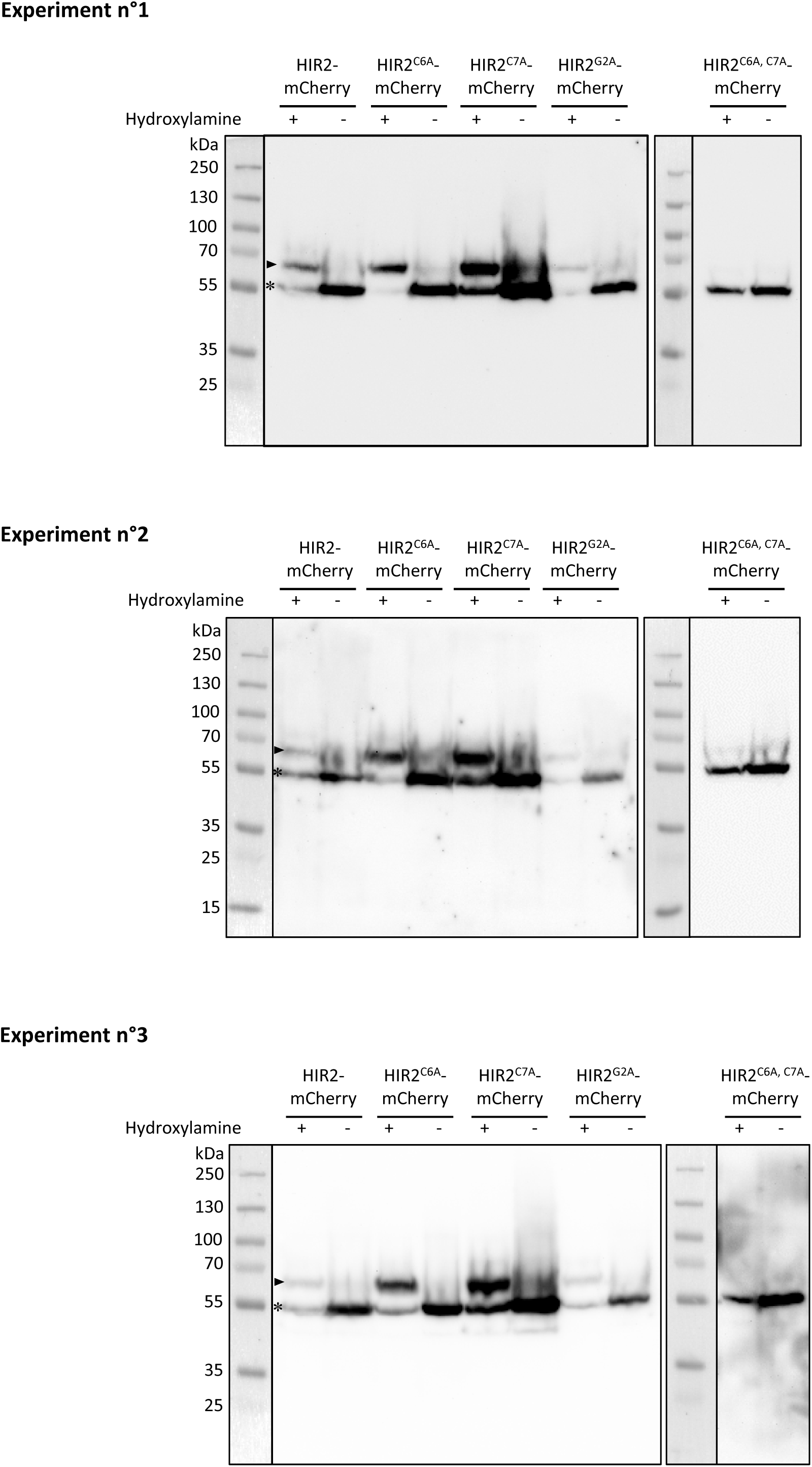
HIR2 is mono S-acylated and simultaneous mutations of C6 and C7 abolish HIR2 S-acylation. The S-acylation of WT HIR2, HIR2^C6A^, HIR2^C7A^, HIR2^G2A^ and HIR2^C6A, C7A^ proteins fused to the mCherry and expressed under the control of *HIR2* promoter in Arabidopsis transgenic lines was assessed with a PEG-shift assay combined with anti-mCherry immunoblots. Following the cleavage of S-acyl groups with the thioester cleavage reagent hydroxylamine, a maleimide-functionalized polyethylene glycol (mPEG-Mal) was selectively coupled to free cysteines, which induced a mobility shift of 10 kDa for each S-acylation site. In the absence of hydroxylamine (-), no mass shift should be observed with a signal at ∼ 55 kDa, corresponding to the expected molecular weight of HIR2 and HIR2 mutated proteins fused to the mCherry. The mono S-acylation of WT HIR2, but also some HIR2 mutated proteins fused to the mCherry is indicated by the detection of a signal at ∼ 65 kDa in the presence of hydroxylamine (+). Arabidopsis seedlings were grown for 11 days on MS/2 medium. Non-S-acylated and mono S-acylated HIR2 and HIR2 mutated proteins fused to the mCherry are indicated by a star and an arrowhead, respectively. Three independent experiments corresponding to three biological replicates are shown. Experiment n°1 corresponds to the immunoblot presented in Figure 3B. The protein ladder transferred on the membranes is shown on the left of the immunoblots.

**Supplemental Figure S6.**
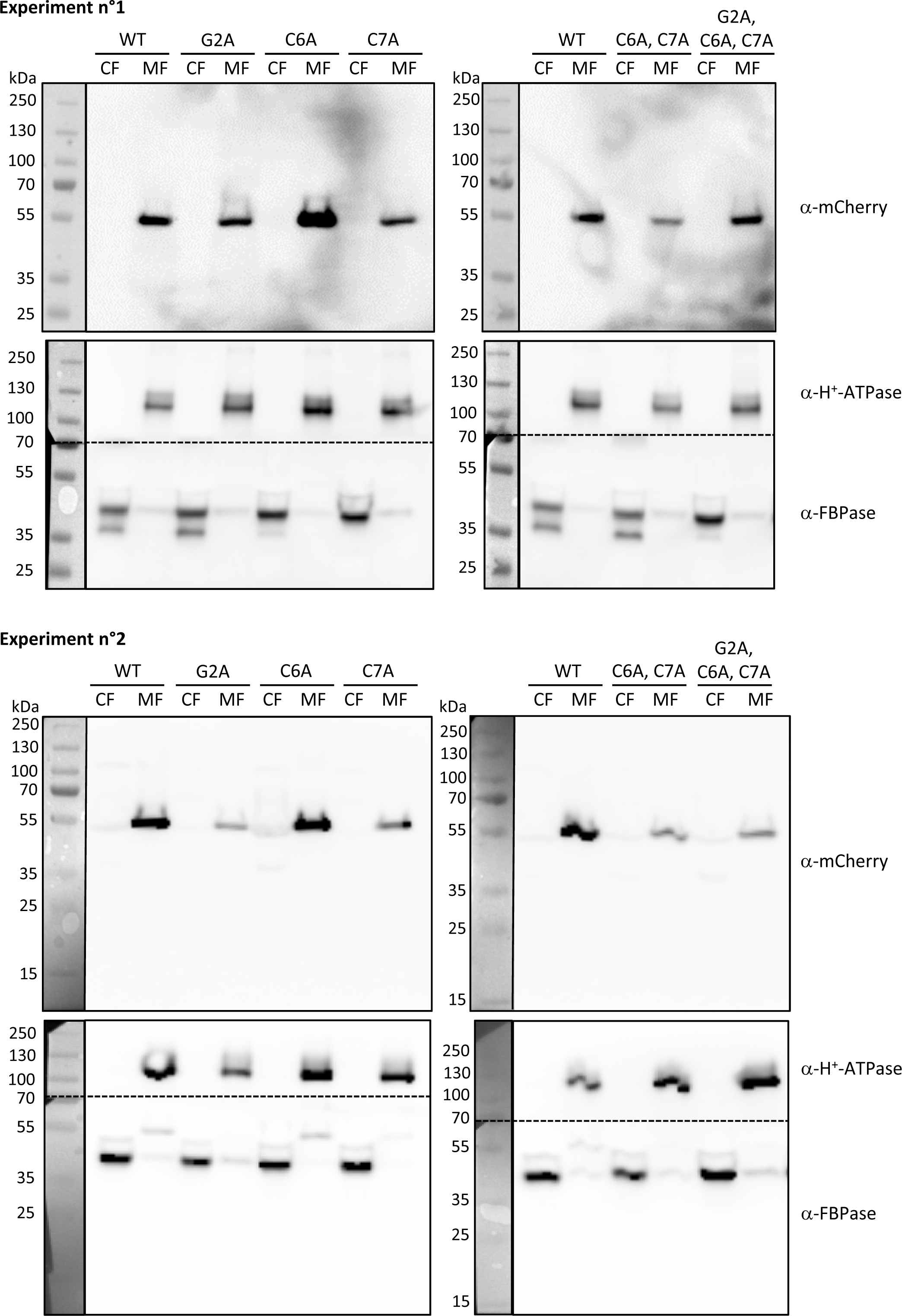
Point-mutated variants of HIR2 affected for S-acylation and myristoylation still associate with membranes. Cell fractionation was performed on *Nicotiana benthamiana* leaves transiently expressing HIR2-mCherry (WT), HIR2^G2A^-mCherry (G2A), HIR2^C6A^-mCherry (C6A), HIR2^C7A^-mCherry (C7A), HIR2^C6A, C7A^-mCherry (C6A, C7A) and HIR2^G2A, C6A, C7A^-mCherry (G2A, C6A, C7A) proteins under the control of *HIR2* promoter. Cytosolic and microsomal protein fractions, noted CF and MF, respectively, were probed with an anti-mCherry antibody. After stripping, the transfer membranes were cut horizontally (dash lines) and re-probed with anti-H^+^-ATPase (upper part), and anti-cytosolic FBPase antibodies (bottom part). AHA and FBPase proteins were used as markers of membrane and cytosolic fractions, respectively. Two independent experiments corresponding to two biological replicates are shown. The protein ladder transferred on the membranes is shown on the left of the immunoblots.

**Supplemental Figure S7.**
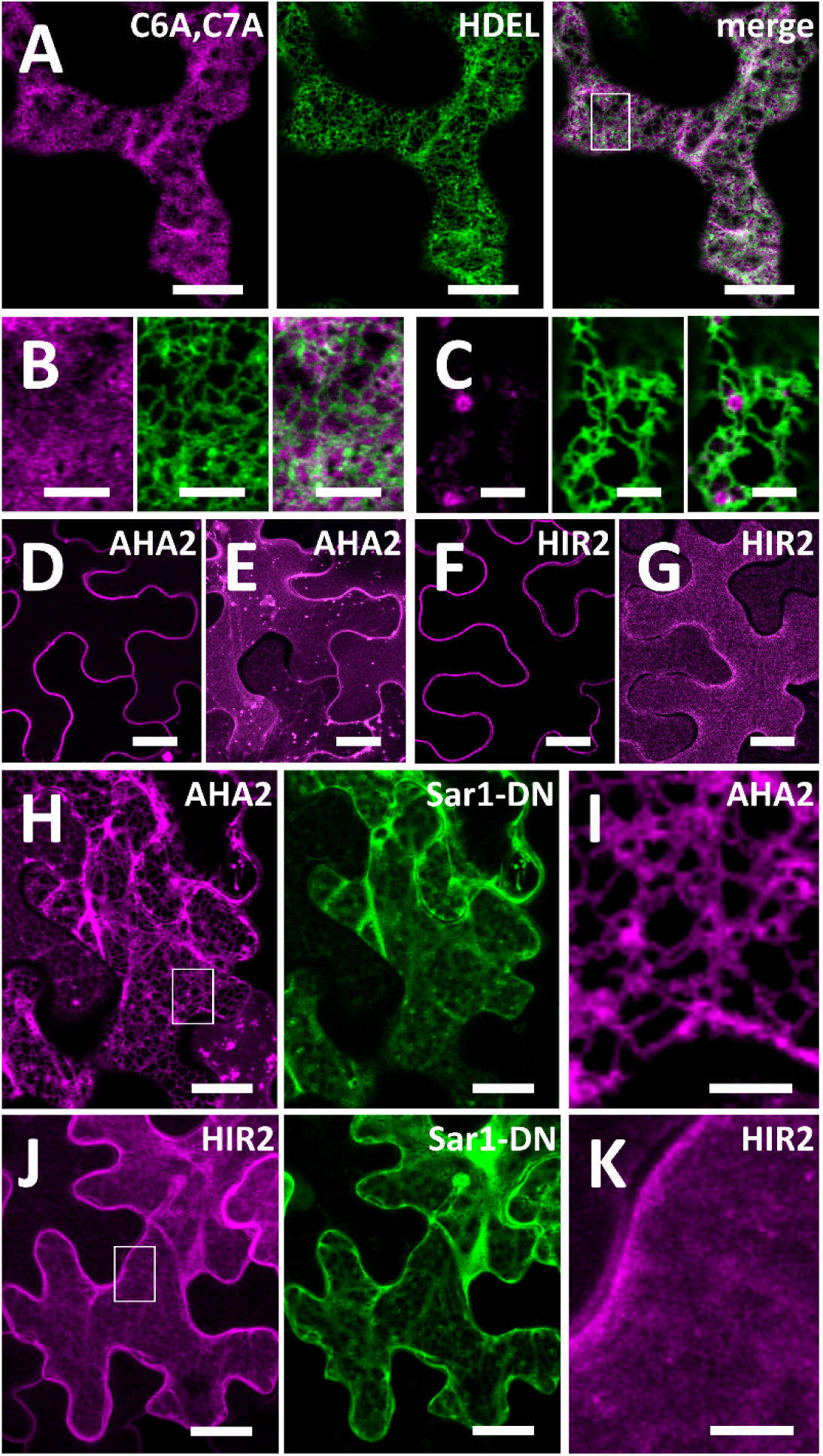
HIR2 plasma membrane targeting relies on mechanisms independent of the secretory pathway. **(A)** The intracellularly-retained HIR2^C6A, C7A^ protein does not co-localize with the ER marker pHluorine-HDEL. HIR2^C6A, C7A^-mCherry (left panel) and pHluorine-HDEL (middle panel) proteins were co-expressed in *Nicotiana benthamiana* leaf epidermal cells using *HIR*2 and *35S* promoters, respectively. Observation were performed by scanning confocal microscopy, maximum intensity projection of a Z-stack covering the cell surface is presented. The superimposed image for the two channels (merge) is shown in the right panel. **(B)** Close-up view of the region marked with a white rectangle in (A). **(C)** HIR2^C6A, C7A^-mCherry protein is occasionally observed in round shape structures (left panel) that do not co-localize with the ER marker pHluorine-HDEL (middle panel) as seen in the merged images (right panel). **(D – K)** Contrary to the H^+^-ATPase AHA2, the trafficking of HIR2 to the PM does not require the COPII machinery. AHA2-mCherry **(D-E)** and HIR2-mCherry fusion proteins **(F-G)** are both localized in the PM when expressed in *N. benthamiana* leaf epidermal cells. **(D)** and **(F)** Secant views. **(E)** and **(G)** Maximum intensity projection of a Z-stack spanning from the cell surface to the position shown in (D) and (F), respectively. Note that AHA2 is also detected in some intracellular vesicles (E). **(H)** AHA2-mCherry is retained in the ER (left panel) when co-expressed with a dominant-negative (DN) form of Sar1 fused to the YFP (right panel) in *N. benthamiana*. **(I)** Close-up view of the region marked with a white rectangle in (H). **(J)** HIR2-mCherry is not retained in the ER and is mostly delivered to the PM (left panel) when co-expressed with Sar1DN-YFP (right panel) in *N. benthamiana*. In (H) and (J), a maximum intensity projection of a Z-stack covering the cell surface is presented. **(K)** Close-up view of the region marked with a white rectangle in (J). Scale bars represent 20 µm (A, D – H, J) and 5 µm (B, C, I, K).

**Supplemental Figure S8.**
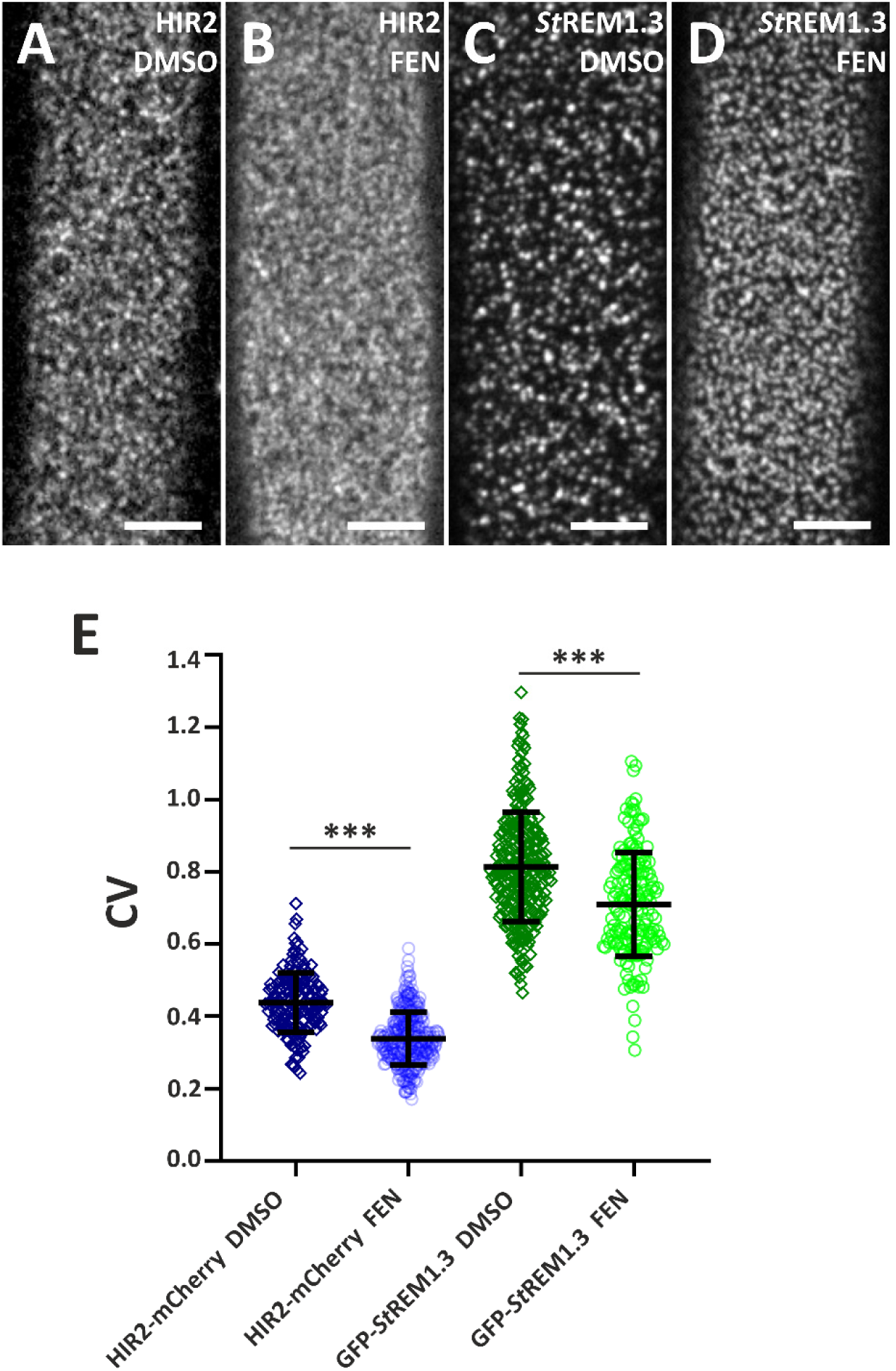
Fenpropimorph treatment alters the nanodomain organization of both HIR2-mCherry and GFP-StREM1.3 proteins at the plasma membrane in Arabidopsis root epidermal cells. (**A-D**): Spinning disk microscopy analysis on the plasma membrane of root epidermal cells from 10-day old Arabidopsis seedlings expressing pHIR2::HIR2mCherry (A, B) or 35S::GFP-StREM1.3 (C, D) and mock treated with DMSO (A, C) or treated with 50 µg / mL fenpropimorph (FEN) (B, D) for 24 hours. Scale bars represent 5 µm. (**E)** Coefficient of variance (CV) of fluorescence intensity were calculated from TIRF experiments performed as in (A-D). Scatter plots with indicated means ± standard deviation are presented. Asterisks indicate significant differences (*** p< 0.001, two-sample t-test, n = 173 – 328 intensity profiles of 8 – 12 seedlings per variant). Three biological replicates providing similar results were performed independently.

**Supplemental Figure S9.**
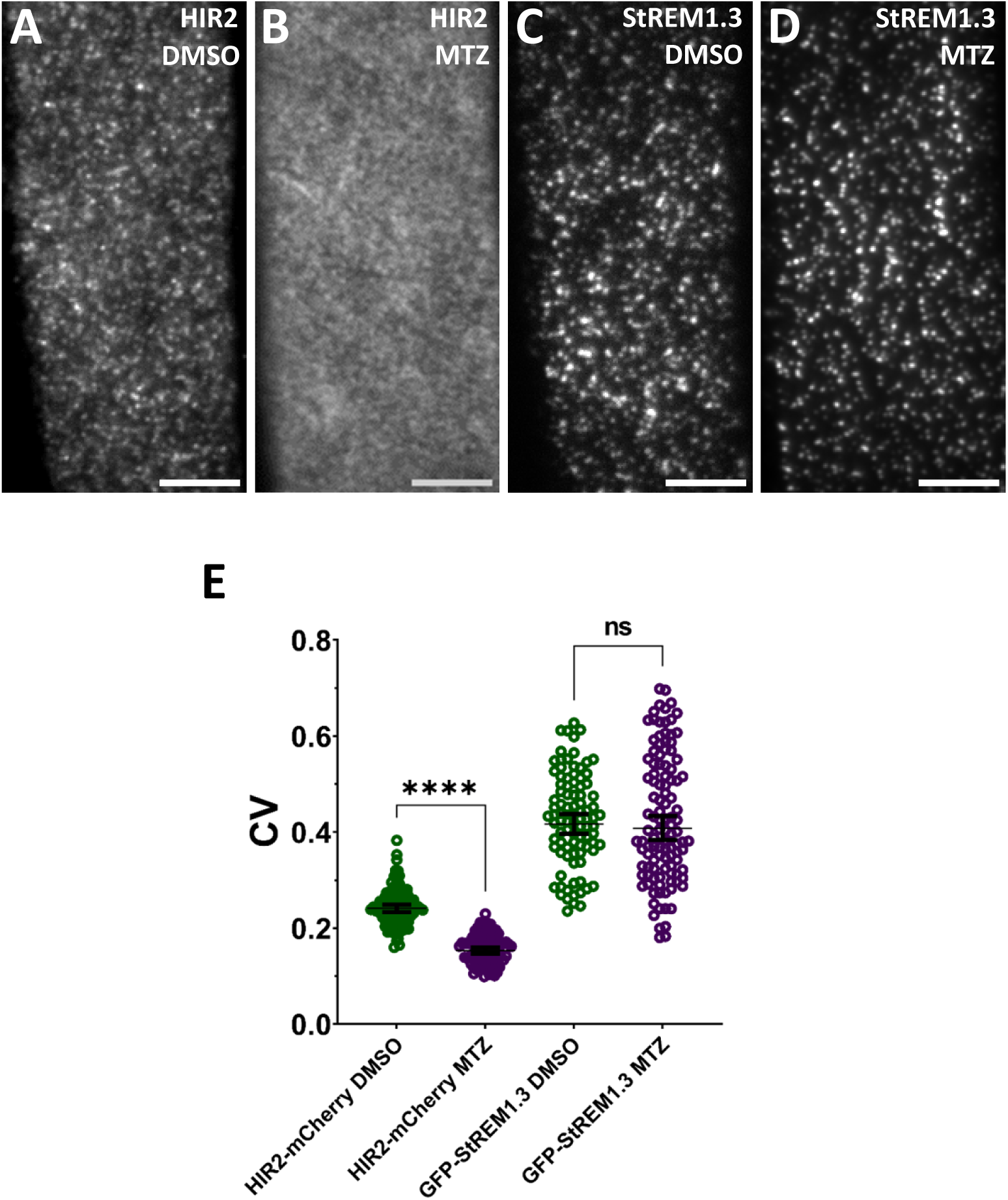
StREM1.3 nanodomain organization is not disturbed by metazachlor. **(A-D)** TIRF microscopy analysis on the PM of root epidermal cells from transgenic Arabidopsis seedlings co-expressing pHIR2::HIR2-mCherry and 35S::GFP-StREM1.3 mock-treated with DMSO (A and C) and treated with metazachlor (MTZ) (B and D). (A, B): HIR2-mCherry fluorescence, (C, D): GFP-StREM1.3 fluorescence. Scale bars represent 5 µm. (**E)** Coefficient of variance (CV) of fluorescence intensity determined from TIRF images of root epidermal cells co-expressing HIR2-mCherry and 35S::GFP-StREM1.3 and treated as in (A-D). Error bars correspond to the mean with a confidence interval at 95%. Asterisks indicate significant differences (**** p< 0.0001, in a t-test), ns means non-significant, n = 20 – 25.

**Supplemental Figure S10.**
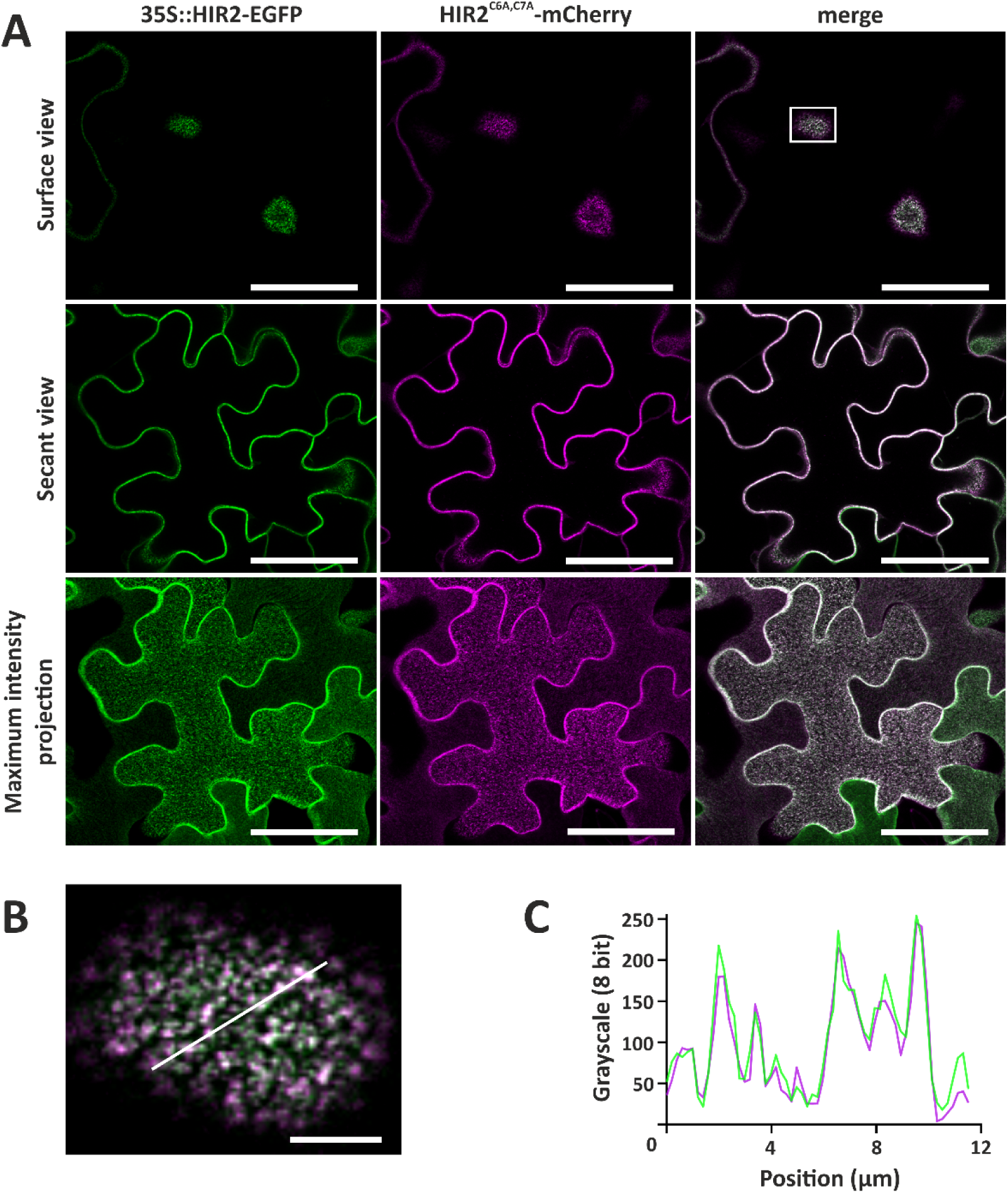
HIR2^C6A, C7A^-mCherry is re-localized to plasma membrane nanodomains upon co-expression with the non-mutated HIR2 protein fused to EGFP. HIR2^C6A, C7A^-mCherry and HIR2-EGFP proteins were co-expressed in *N. benthamiana* leaf epidermal cells using *HIR2* and the *35S* promoters, respectively. Observation were performed by scanning confocal microscopy. **(A)** HIR2-EGFP (left panel) and HIR2^C6A, C7A^-mCherry (middle panel) are localized in the PM, where they form clusters that co-localize (merge, right panel). Upper panels: surface view at the top of the cell. Middle panels: secant view. Lower panels: maximum intensity projections of a Z-stack covering the cell surface. **(B)** Close-up view of the region marked with a white rectangle in (A). (**C)** Signal intensity profiles of 8-bit images for EGFP channel (green line) and mCherry channel (magenta line) obtained from the linear ROIs (white line) in (B). Note the overlap of the signals illustrating the co-localization of both proteins within PM nanodomains. Scale bars: 50 µm (A) and 5 µm (B).

**Supplemental Figure S11.**
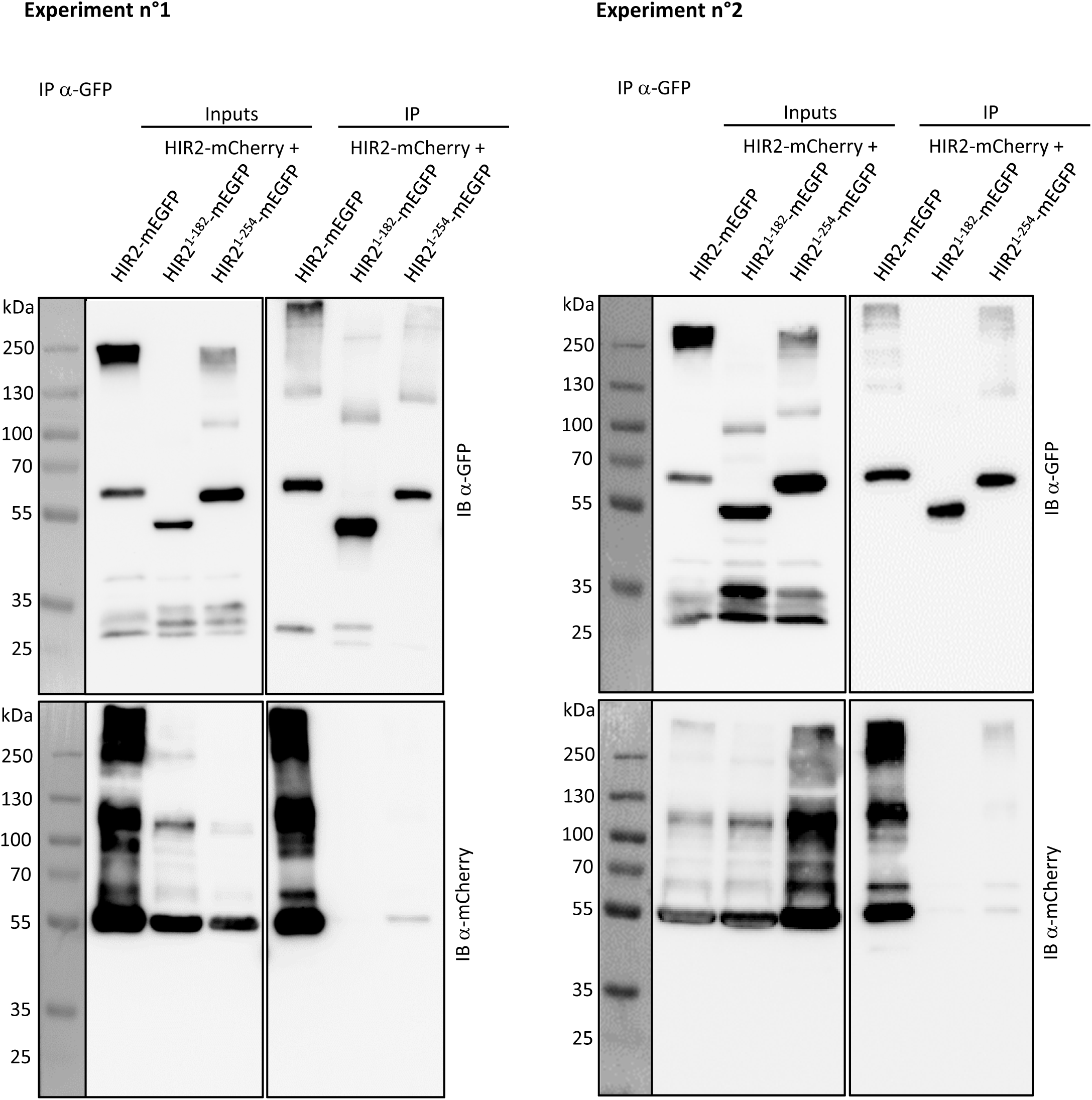
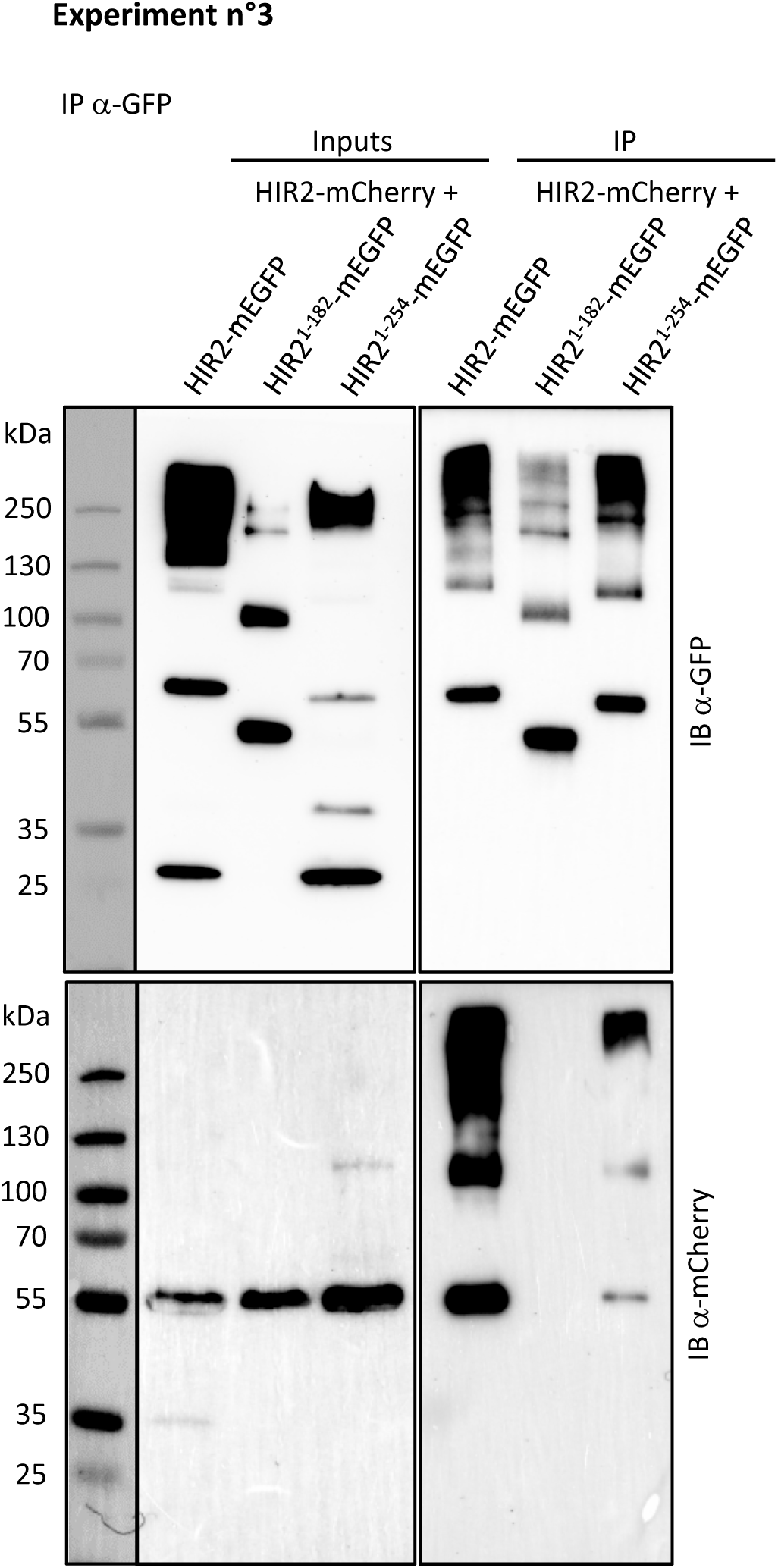
The C-terminal part of HIR2 is required for oligomerization as revealed by co-immunopurifications. Immunopurifications (IP) using an anti-GFP antibody were performed on solubilized protein extracts from *N. benthamiana* leaves transiently co-expressing HIR2-mCherry with HIR2-mEGFP, HIR2^1-182^-mEGFP or HIR2^1-254^-mEGFP. Inputs and IP fractions were subjected to immunoblots (IB) with anti-GFP (top) and anti-mCherry antibodies (bottom). The expected molecular weights of HIR2-mCherry, HIR2-mEGFP, HIR2^1-182^-mEGFP and HIR2^1-254^-mEGFP proteins are 58.9 kDa, 59.3 kDa, 48.6 kDa and 56.3 kDa, respectively. Some signals can be observed for higher and lower molecular weights and probably correspond to not fully denatured and cleaved fusion proteins, respectively. Three independent experiments corresponding to three biological replicates are shown. Experiment n°1 corresponds to the immunoblots presented in Figure 6J. The protein ladder transferred on the membranes is shown on the left of the immunoblots.

**Supplemental Figure S12.**
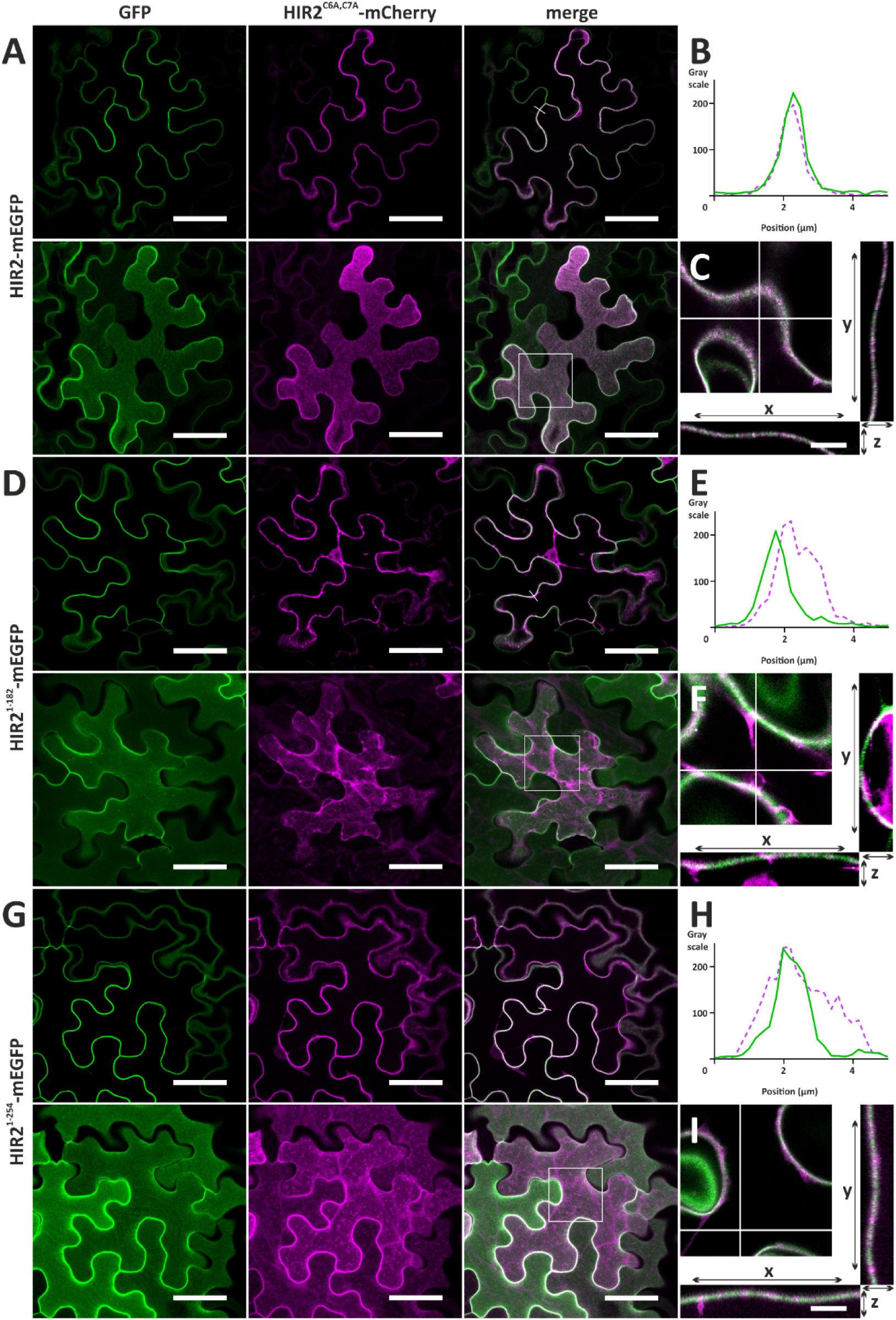
Plasma membrane re-localization of HIR2^C6A, C7A^-mCherry is ineffective upon co-expression with C-terminal deleted versions of HIR2. HIR2^C6A, C7A^-mCherry protein was co-expressed with HIR2-mEGFP **(A)**, HIR2^1-182^-mEGFP **(D)** and HIR2^1-254^-mEGFP **(G)** in *N. benthamiana* leaf epidermal cells using *HIR2* promoter. HIR2^1-182^ and HIR2^1-254^ are truncated proteins corresponding to the first 182 and 254 amino acids of HIR2, respectively. Observation were performed by scanning confocal microscopy. **(A, D, G)**: Upper panels: secant view; lower panels: maximum intensity projections constructed from Z-stack of 25 focal planes spanning from the epidermis surface to the positions shown in upper panels. (**B, E, H)** Signal intensity profiles of 8-bit images for EGFP (full green line) and mCherry (dashed magenta line) obtained from the linear ROIs (white lines) shown in upper merge images of (A), (D) and (G). Note the almost complete overlap of the peaks in (B), shift of the peaks in (E) and broader peak corresponding to mCherry signal that partially co-localize with mEGFP peak in (H). (**C, F, I)** XZ and YZ orthogonal projections from ROIs (white squares in lower merge images) of the Z-stacks corresponding to (A), (D), (G). The positions of the projections in XY is marked with white crosses in the insets. Scale bars represent 50 µm (A, D, G), and 10 µm (C, F, I).

**Supplemental Table 1. Primers used in this work.**

**Supplemental Video 1. spt-PALM analysis of the dynamics of HIR2-mEOS2 single molecules in the plasma membrane of Arabidopsis root epidermal cells.** Arabidopsis seedlings stably expressing HIR2-mEOS2 fusion protein under the control of *HIR2* promoter were imaged.

## Notes

### Competing Interest Statement

The authors have declared no competing interest.

